# A wireless, user-friendly, and unattended robotic flower system to assess pollinator foraging behaviour

**DOI:** 10.1101/2022.06.14.496104

**Authors:** Kamiel Debeuckelaere, Dirk Janssens, Estefanía Serral Asensio, Tom Wenseleers, Hans Jacquemyn, María I. Pozo

**Affiliations:** Laboratory of Socioecology and Social Evolution, KU Leuven, Naamsestraat 59 - box 2466, 3000 Leuven, Belgium; Plant Population Biology and Conservation, KU Leuven, Kasteelpark Arenberg 31 - box 2435, 3001 Leuven, Belgium; ICTS - Central Infrastructure, KU Leuven, Willem de Croylaan 52a - box 5580, 3001 Leuven, Belgium; Research Centre for Information Systems Engineering (LIRIS), KU Leuven, Warmoesberg 26 - box 15101, 1000 Brussels, Belgium

**Keywords:** automated data collection, conservation ecology, floral rewards, insect behaviour, Internet of Things (IoT) applications, pollination ecology, wireless sensor networks

## Abstract

1. Understanding the complex interactions between external and internal factors that influence pollinator foraging behaviour is essential for developing effective conservation strategies. However, collecting large datasets that incorporate data from various sources has been challenging.
2. To address this issue, we present a wireless and cost-effective robotic flower equipped with Internet of Things (IoT) technology that automatically offers nectar to visiting insects while monitoring visitation time and duration. The robotic flower is easy to manipulate and settings such as nectar refill rates can be remotely altered, making it ideal for field settings. The system transmits data completely wirelessly and autonomously, is mobile and easy to clean.
3. The prototype settings allow for approximately two weeks of uninterrupted data collection for each battery charge. As a proof-of-concept application, a foraging-preference dual choice experiment with bumblebees was performed. On average, more than 14 000 flower visits were registered daily with a setup consisting of 16 robotic flowers. The data show a gradual preference shift from the pre-trained, lower quality food source towards the higher quality source.
4. The robotic flower provides accurate and reliable data on insect behaviour, dramatically reducing the price and/or labour costs. Although primarily designed for (bumble)bees, the system could be easily adapted for other flower-visiting insects. The robotic flower is user-friendly and can be easily adapted to address a wide range of research questions in pollination ecology, conservation biology, biocontrol and ecotoxicology, and allows for detailed studies on how nectar traits, flower colour and shape and pollutants would affect foraging behaviour.

## Introduction

“*If insects were to vanish, the environment would collapse into chaos*.” - E.O. Wilson

Pollinators play a crucial role in the sexual reproduction of approximately 90% of all flowering plants (Ollerton et al., 2011), including many species of agricultural importance (Klein et al., 2007). Among pollinators, insects and especially bees play a key role (Potts et al., 2010). The transfer of pollen from female to male plant organs is mediated by complex interactions between plants and insects and largely depends on the foraging behaviour of individual insects, which in turn depends on both internal and external factors. While flower handling abilities, foraging experience, reward recognition capabilities, and insect or colony health are the main internal factors affecting pollinators’s foraging decisions (Lunau, 2000; Lunau et al., 1996; Minahan & Brunet, 2018; Schmid-hempel & Stauffer, 1998; Spaethe & Weidenmüller, 2002), maximization of energy gain through the collection of floral rewards is also affected by abiotic features such as light and temperature, landscape features, flower density, colour and shape of the flowers, and the quantity and quality of floral rewards (Nicolson, 2011; Spaethe et al., 2001; Sun et al., 2018; Vicens & Bosch, 2000; Waddington & Holden, 1979; Werner et al., 2016). The relative importance of these factors and how they interact with each other is not yet completely understood, especially for non-model organisms (e.g., flies (Hannah et al., 2019)).

Behavioural experiments with pollinators have proven to be challenging as they often imply collecting data from fast moving animals and multiple individuals simultaneously (Dankert et al., 2009; Dell et al., 2014). Manual collection of such behavioural data relies on continuous observation and is thus labour intensive, time consuming and subject to human limitations in observing dynamic objects (Crall et al., 2015; Simons & Chabris, 1999). In addition, invasive human actions are often needed, which could potentially influence foraging behaviour (Calisi & Bentley, 2009). To counter this, researchers often used simplified devices to perform manipulative research, thus being able to address complex biological questions in a systematic way (see, e.g., Goulson and Cory (1993); Gumbert (2000); Real (1981)). Alternative monitoring strategies have been developed as well (see Table 1 for examples), but these often require specialized, complex and expensive equipment. Cheap, reliable, automated, and easy to use methods can help to enhance the quality and quantity of data on foraging behaviour in pollinators.

**Table 1:**
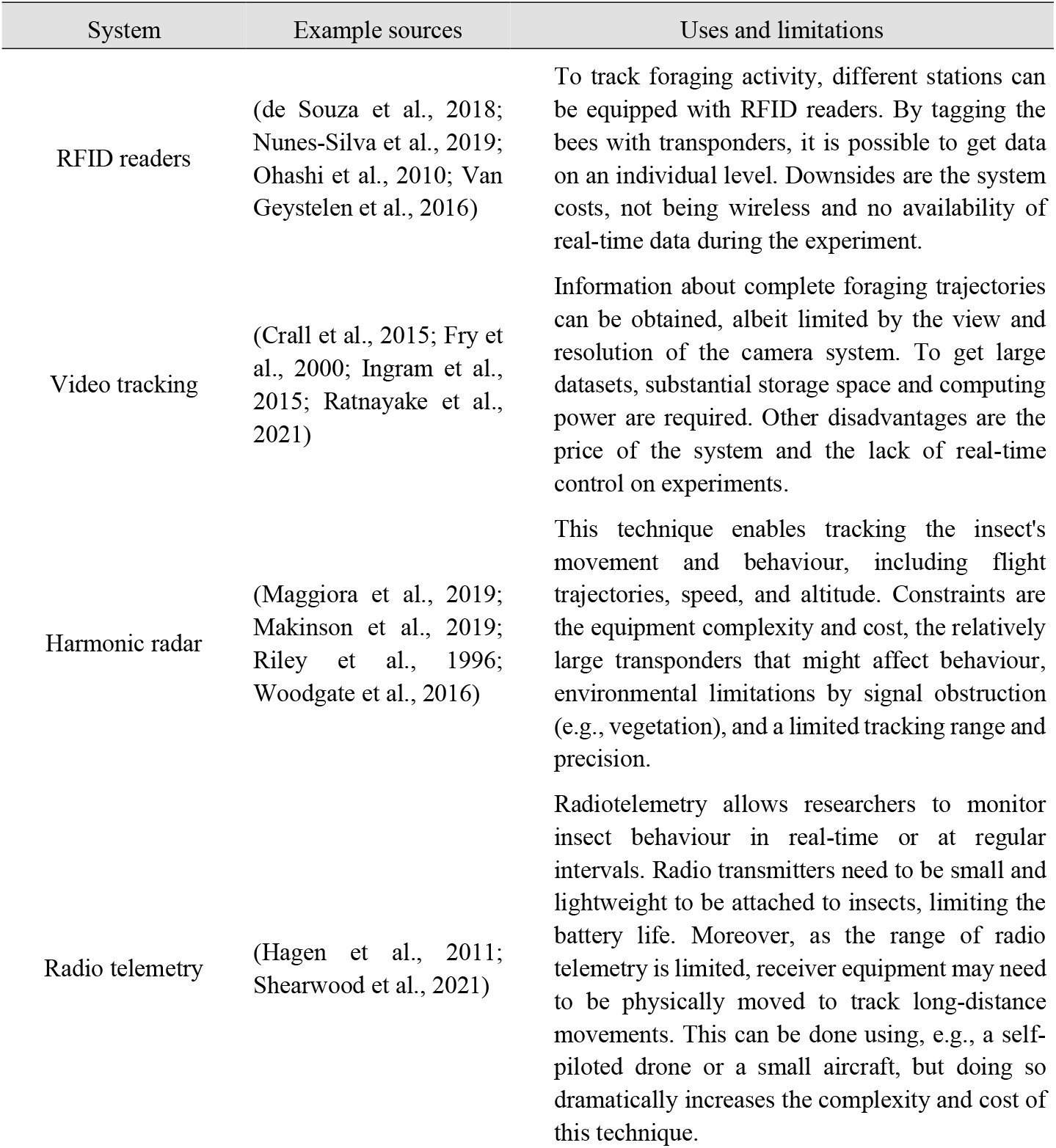
Overview of some other foraging behaviour monitoring strategies.

Robotic flowers, which have mechanical features and can function without the requirement of human intervention, are regarded as a type of artificial flowers. In the past years, various robotic flower systems have been constructed with many different mechanisms of reward delivery (Essenberg, 2015). Although non-nectar rewarding systems have been made (see, e.g., Switzer et al. (2019)), most robotic flowers offer nectar, which is the main reward of plants for their mutualistic counterparts (Nicolson & Thornburg, 2007). Nectar can be delivered in continuous doses (Ings & Chittka, 2008; Makino & Sakai, 2007; Moffatt, 2001; Ohashi et al., 2010; Paldi et al., 2003; Thomson et al., 2012) or discrete doses (Cnaani et al., 2006; Essenberg, 2015; Greggers & Menzel, 1993; Keasar, 2000; Nachev & Winter, 2012; Sokolowski & Abramson, 2010). The ability to automatically detect a floral visitor was also implemented in some systems, using an infrared detection system (Greggers & Menzel, 1993; Keasar, 2000; Kuusela & Lämsä, 2016; Nachev & Winter, 2012). Kuusela and Lämsä (2016) combined the designs noted above and developed a low-cost robotic flower system with automatic visit detection that offers nectar in discrete doses using a servomotor. However, none of these robotic flower systems mentioned above can be used wirelessly. The need to be connected with cables to a central control unit (e.g., a computer) and a power supply makes them less versatile. Fixed systems are often less user-friendly (e.g., with respect to cleaning or fixing damaged parts) and less widely deployable compared to autonomous wireless systems, which can be moved around freely.

In this manuscript, we introduce an updated robotic flower version achieving full wireless functionality through the integration of a battery and a low power, long range wireless data transmission system operating within the Internet of Things (IoT) framework. In order to optimize the potential duration of experiments using a single battery, significant emphasis was placed on enhancing the power efficiency of the system. The robotic flower system developed here was specifically designed for (bumble)bees, but with minor adjustments (e.g. of the size of the feeding hole and/or nectar cup) larger or smaller floral visitors, such as syrphid flies, could be investigated as well.

## Description and implementation

The subsequent paragraphs will provide a comprehensive overview of our innovative robotic flower. Firstly, the system architecture is clarified, elucidating the underlying technology. Subsequently, we will delve into the intricate design of the robotic flower, explaining the materials employed and the mechanisms incorporated in its construction. Lastly, we will present a proof of concept experiment, showcasing the efficacy of the robotic flower and its potential applications.

### System Architecture

The system consists of separately operating robotic flowers that register the time and duration of every visit via an infrared sensor and transfer these data wirelessly (IoT) to be stored in a CSV-file (Fig. 1). The user can remotely select and change the nectar refilling rate through a dashboard website that also allows monitoring (e.g., the battery level) of all the robotic flowers in use (Fig. S1). The time-frame for a sleep mode to save battery power can also be selected here.

**Figure 1:**
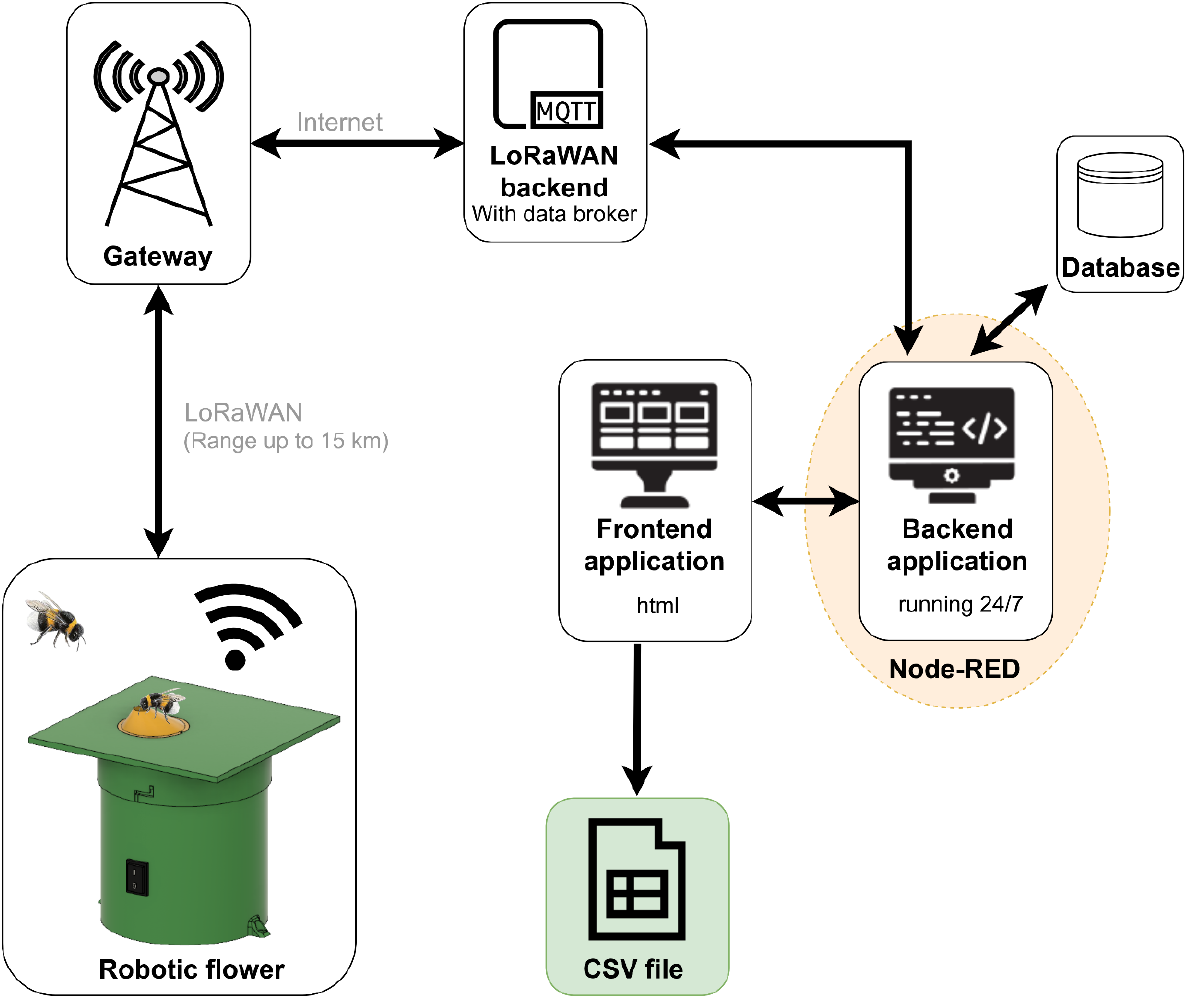
Overview of the robotic flower system using IoT technology. The robotic flower sends data packages using LoRaWAN (Long Range Wide Area Network) to a gateway, which forwards the message through internet to the LoRaWAN-backend. Here, the MQTT (message queue telemetry transport) broker acts like a post office: it receives all messages sent from devices (publishers) and then forwards it to the right applications (subscribers). These messages contain two parts: the topic and the payload. The topic identifies the message, while the payload contains the visitation data (time and duration of each visit), battery level, alarm status, and metadata (e.g., the time of sending, a list of gateways that received the transmitted message and the speed of data transmission). The application has a backend that is constantly running to receive this message and store data in the database, and a frontend that enables the user to overlook all deployed robotic flowers in a dashboard website. The visitation data transmitted by the robotic flowers are gathered in daily CSV files. After receiving a message, the application sends a reply back to the robotic flower in the same way but in opposite direction. This message contains a receive confirmation and instructions about sleep mode and nectar refilling rate.

Each robotic flower has its own Arduino microcontroller and a transceiver based on wireless LoRa technology for communication with LoRaWAN (Long Range Wide Area Network). LoRaWAN is a low power networking protocol used for IoT applications in which a single gateway can receive data from thousands of devices in a radius of 2 to 5 km in urban areas, and up to 15 km in more remote areas (Adelantado et al., 2017). The gateway is listening to all incoming LoRa signals and translates the incoming radio waves into electrical signals. The gateway, which is connected to the internet, then forwards the received message from devices (publishers) to the right applications (subscribers) through a LoRaWAN backend server with an MQTT (message queue telemetry transport) broker (Provoost & Weyns, 2019). This project made use of DingNet, the LoRaWAN communication infrastructure of KU Leuven, which provides the LoRaWAN backend and the gateways with antennas that ensure coverage across the whole city (Provoost & Weyns, 2019). The Things Network (The Things Industries B.V.), which provides a free LoRaWAN backend globally, is an example of an alternative infrastructure that could be used.

The robotic flower publishes messages with a regular interval (10 minutes in the prototype) with a maximum size of 51 bytes each. This is to limit the consumed airtime and ensure fair spectrum division with other users (Adelantado et al., 2017). In the case where more visits are registered than can be send at once, data is temporarily stored at the back of the sending queue, waiting for the next transmission. In this way, data for more than 1500 visits per day per flower can be stored in the CSV-file.

The application to receive the LoRaWAN messages was developed with Node-RED (OpenJS Foundation). To ensure continuous operation of Node-RED while the robotic flowers are in use, a computer (localhost) was employed, although alternative options are also available, e.g., with a Raspberry Pi, with a server or in the cloud. The application stores the visitation data in a daily CSV-file: for every visit, a new line with the time at the start of the visit, the duration of the visit, and the name of the device that registered the visit (e.g., ‘flower 1’) is added to the file. When the application receives a message from the robotic flower, a small message is sent back as an answer to the flower containing the currently selected nectar refilling rate, the order to go to sleep or wake-up, and to confirm whether the data transmission was successful or not. If unsuccessful, the flower will try to re-send the data.

The firmware, written in C++, runs on the Arduino microcontroller of each flower. When this microcontroller is powered on, e.g., by switching on the battery power with the rocker switch, the program will start running. This always starts with a setup loop to initialize the robotic flower functions and establish the LoRaWAN connection before continuing with the main loop, which keeps running as long as power is supplied. The flower is programmed to have two states: working and sleeping. In the working state, the infrared detection system is functional and data is regularly send. These functions can be stopped during sleep to save battery life when the targeted study species are not active during the night (e.g., bumblebees (Spaethe & Weidenmüller, 2002)). The firmware also allows adjusting the hardcoded behaviour of the flower. To do so, the updated firmware needs to be re-uploaded with a USB-cable to the microcontroller. Specifically, the following variables can be easily adjusted: period of deep sleep, sensor sensitivity (i.e., how big the voltage drop in the infrared sensor needs to be to detect a visitor), automatic nectar refilling frequency (to prevent drying out), different options for the nectar refill-gap duration, data transmission frequency, maximum visit duration, battery voltage warning threshold, visit duration threshold to trigger a nectar refilling, and the threshold to give an obstruction alarm.

### Robotic Flower Design

The mechanical design of the robotic flower is based on the previous design by (Kuusela & Lämsä, 2016). A visit is recorded by a voltage drop of the infrared sensor when it is blocked by a bumblebee in the feeding hole (Fig. 2). This detection system has been used before and proved to be robust, straightforward, and reliable (Keasar, 2000; Lämsä et al., 2018; Sokolowski & Abramson, 2010). Nectar is offered in discrete amounts to simulate the natural availability of nectar in flowers and to stimulate foraging in floral patches. The refilling of artificial nectar from the nectar reservoir is done with a servo-arm and a small nectar cup connected to a servomotor (Fig. 2).

**Figure 2:**
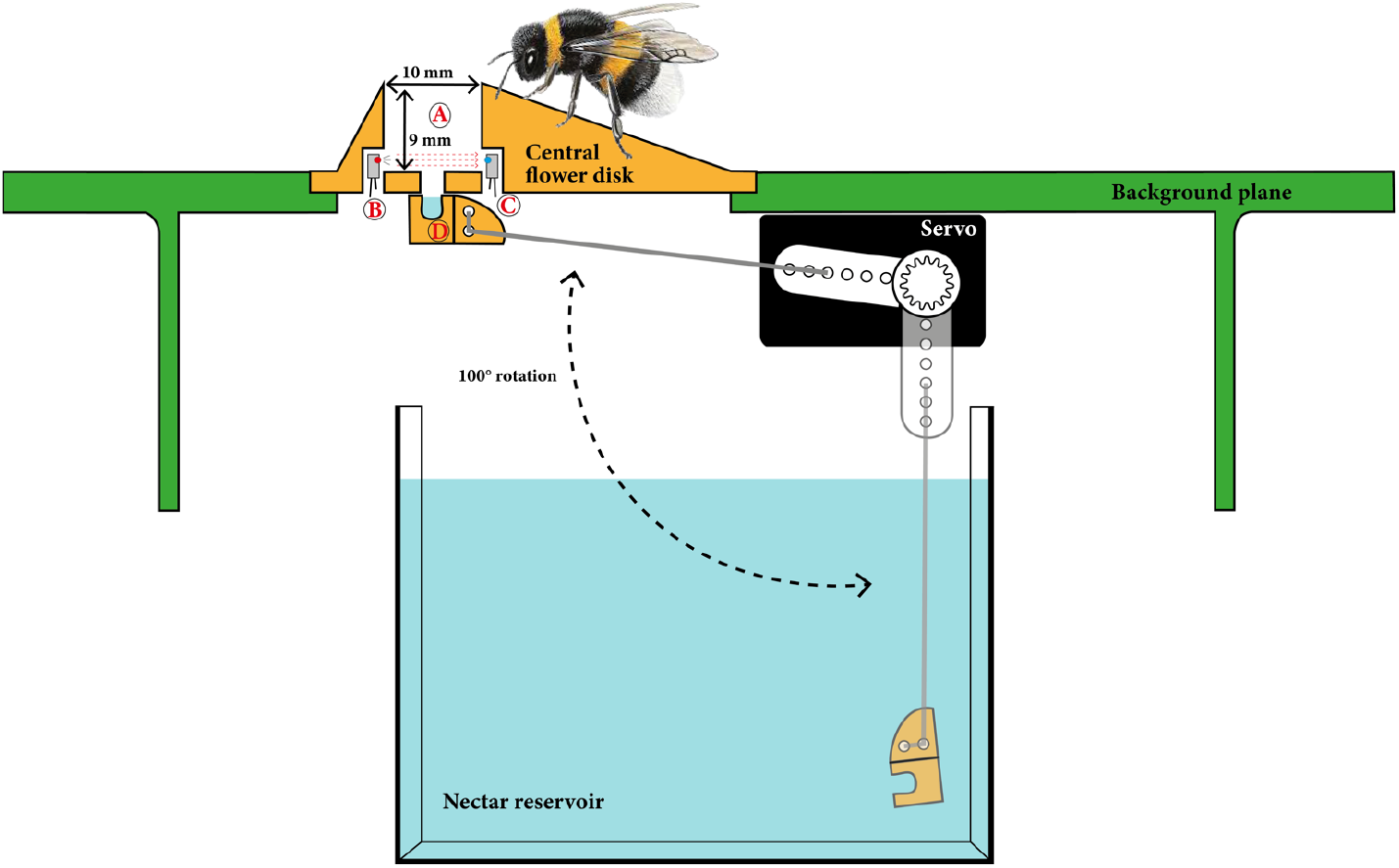
Mechanical design of the robotic flower based on Kuusela & Lämsä (2016) with A) the feeding hole, B) the infrared emitter, C) the infrared sensor, and D) the nectar cup. The servo-arm is used to refill the nectar cup from the nectar reservoir (rotation of 100°) and press it to the central flower disk, where floral visitors can reach the nectar using their proboscis.

The design of the robotic flower was made using Autodesk Fusion 360. Each flower consists of five 3D-printed components that fit together: stem, background plane, central flower disk, nectar cup, and reservoir case (see part ‘Data accessibility’ for the design files). The used 3D-printer was a Prusa I3 MK3S (Prusa Research a.s., Prague), with a mixture of PLA (polylactic acid) and colourants as printing material. An animation of the robotic flower assembly and detailed pictures with information about the design and construction can be found in the supporting information (video S2 and file S3). All parts can be disassembled easily to facilitate cleaning, and electronics are protected either from internal spills or external dirt. The weight of the total system is approximately 370 g.

The stem serves as the housing part for the electronic components of the robotic flower. In this way, internal parts are hidden so that foraging behaviour cannot be influenced by, for example, observable motion of the servo-arm. The background plane is shaped as a square, making it possible to form one continuous platform when grouping multiple flowers together (Fig. S5). The colour green was chosen to make a cryptic background (Chittka et al., 1994). The central flower disk represents the actual flower and was designed with a wide diameter (45 mm) and colour contrasting the green background (bright yellow and blue were selected for the prototype) so that detectability by bees is enhanced during flight (Spaethe et al., 2001). The central flower disk contains a feeding hole with an infrared detection system designed specifically for medium/large bees (> 10 mm long), which covers the most relevant pollinators in temperate areas such as honeybees, bumblebees, mason bees, and leaf cutting bees (Goulson (2003); Fig. 2). At the bottom of the feeding hole, there is a nectar hole through which the nectar cup can be probed. Transparent silicone is used to attach the central flower disk to the background plane.

The volume of the nectar cup can be altered to meet specific research requirements by changing the 3D-design. Under natural conditions, the volume of nectar has substantial interspecific and intraspecific variances (Real & Rathcke, 1988). In this prototype, the robotic flower presents 15.73 ± 2.30 µL of artificial nectar to visitors in the central flower disk (measured with a 30% sugar solution; see supporting file S6). This discrete volume is not enough to let a visiting bumblebee fill its honey stomach (according to Goulson (2010), ranging between 60 and 120 µL, depending on the size of the worker) with only one visit. Therefore, it is expected to stimulate pollinator movement among the different robotic flowers. The nectar cup is replenished with nectar at a basic refill rate (every 10 min. in the prototype firmware) if visits do not occur. This is to prevent evaporation of the nectar over time. On the other hand, when a visit to the robotic flower reaches a certain threshold duration (e.g., a visit lasting at least 3 sec.), a nectar refill will occur after the visit ends. However, the robotic flower will not refill immediately, but rather leave a lag. This waiting period was installed to match the deterrent action of scent marks on visited flowers (Cnaani et al., 2006). The user can select one of the predefined options for the refill-gap duration from the dashboard website for each individual robotic flower, which can also be changed during an experiment. The refilling itself takes approximately three seconds to avoid formation of air bubbles in the nectar cup.

The reservoir case is built to carry the nectar reservoir while also forming a protective barrier between the electronic components at the bottom and the artificial nectar located at the top. The nectar reservoir itself holds the required artificial nectar for the specific experimental setup. For this prototype, an off-the-shelf reservoir with a maximum volume of approximately 80 ml was used.

Finally, the internal hardware of the robotic flower consists of different electronic parts, of which many are soldered onto a printed circuit board (PCB). A separate battery module is added to use the battery. Hardware parts were selected so they are easy to solder manually. The supporting information contains an overview of the electrical schematic (Fig. S7) and a list of all the parts used, where they were bought, and their article number (supporting file S8). The Gerber files that are needed for PCB fabrication can be found in the GitHub repository (see part ‘Data accessibility’).

A rough price estimate of one robotic flower is € 120 (see Table 2). Although, this price can probably be reduced by using cheaper part alternatives.

**Table 2:**
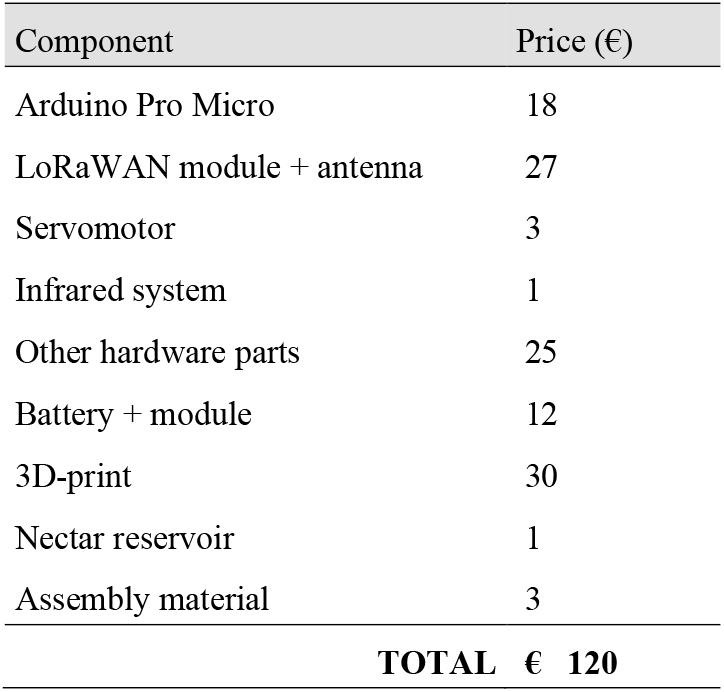
Rough price estimate for parts to build one complete robotic flower (transport costs not included).

The expected battery lifetime was tested to be 14 days (see supporting file S3). If longer experimental periods are desired, empty batteries can be easily replaced during a trial.

### Proof of concept experiments

In a first test, the system was used under both controlled greenhouse and natural outdoor conditions to assess 1) the attractiveness of the design, 2) the detectability of different visitors and 3) the accuracy of visit timing. In the open field setting, being a small botanical garden, robotic flower visitors were observed one day in early summer. Different floral visiting species (e.g., solitary bees, syrphid flies, bumblebees, and honeybees) were attracted to the robotic flower and consumed nectar. A voltage drop in the system was obtained for all these visitors with different body sizes, but we recommend calibrating the sensor sensitivity carefully prior to an experiment. The controlled greenhouse setting was used to test the accuracy of visit detection and timing. Commercially available floral visitors from Biobest (Westerlo, Belgium) were placed inside a small flight together with one robotic flower. Two species were tested: *Eupeodes corollae* (approximately 5 mm in length) and *Bombus terrestris* (approximately 20 mm in length). Floral visits were registered manually in order to verify the reliability and precision of the robotic flower. Following a training period, both species exhibited notable interest in the robotic flower, occasionally posing challenges in visually tracking and accurately timing their visits. Furthermore, slight variations in initiating or ceasing the timer, along with potential rounding errors made by the observer, significantly influenced the recorded visit durations, particularly in cases of short visits. Thus, it was concluded that the utilization of the robotic flowers conferred distinct advantages over manual censuses.

To further prove the value of this new robotic flower, we performed a bumblebee foraging-preference experiment with two reward qualities in a greenhouse setting. After training on less rewarding flowers, we expect an initial preference for these lower-quality flowers (flower constancy) with a gradual shift towards the higher quality option during the trial (Heinrich, 1981). To test this, a robotic flower field consisting of 16 robotic flowers, alternating less and more rewarding flowers in a 4×4 setup, was used in two subsequent replicate trials (Fig. 3A). The two types of flowers are represented by different colours, yellow and blue, so that foragers can discriminate between them visually and associate them with reward quality (Hammer & Menzel, 1995). We used Biogluc 30% *w/w* (less rewarding) and 60% *w/w* (more rewarding) as artificial nectar with preservatives to reduce microbial growth (Billiet et al., 2016). For both trials, a 7-weeks old lab-reared *B. terrestris* colony was placed in the arena with the robotic flowers. Before a trial, colonies were fed pollen and Biogluc ad libitum directly in the nest, but Biogluc supply was ceased 24 hours in advance to stimulate foraging. A trial started with 36 hours of training, where the colony was placed in a closed bug dorm with two 30% w/w rewarding flowers (without refilling gap). Subsequently, the bug dorm was opened at night to allow foraging in the arena starting the next morning (Fig. 3B). A refill gap of 30 minutes was implemented during this experimental phase.

**Figure 3:**
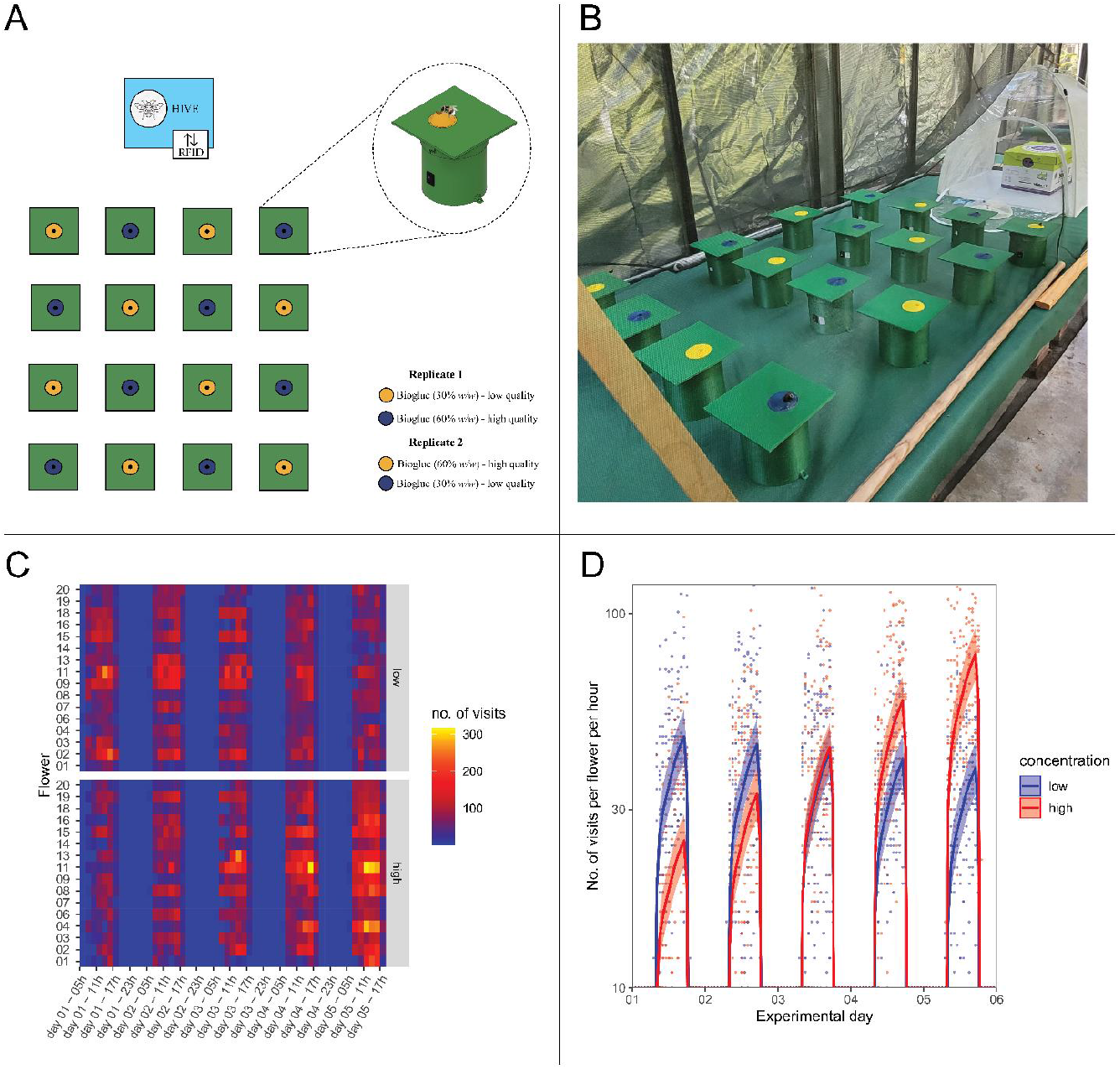
A) Schematical setup of the proof of concept, where in both replicates the low and high quality sources were switched between the yellow and blue flower. B) Picture of the setup, with the bumblebee colony placed in a Bugdorm visible in the back. C) A heatmap of the number of flower visits per quality of the nectar reward over the experimental period. D) An effect plot of the number of bumblebee flower visits per concentration as predicted by the generalized linear mixed model plotted over the raw data. In the first day after being trained on the low concentration, bumblebees visit the low concentration flowers about double as much as they visit high concentration flowers (high/low contrast ratio = 0.527, p < 0.001). There is a linear trend of increasingly more visits towards the high concentration compared to the low concentration (high/low contrast estimate = 0.335, p < 0.001), resulting in no significant difference in visitation rate between the two concentrations on the third experimental day (high/low contrast ratio = 1.030, p = 0.31). On the last experimental day, the preference has switched to little over double the amount of visits to high concentration flowers compared to the low concentration flowers (high/low contrast ratio = 2.015, p < 0.001).

In total, 70 177 visits were recorded in this proof of concept (see CSV file S9). During the daily inspection of the setup within peak activity time-slots, a maximum of five foragers at once were observed. with the earliest flower visit at 7:36 and the latest at 19:17, the natural light regime of October in Leuven (sunrise ∼7:55, sunset ∼19:00) was well visible. The data were analysed using R version 4.2.1 (R Core Team, 2022). A heatmap representing the number of visits per flower per treatment (Fig. 3C) already indicates an initial preference for the low concentration flowers with a switch towards the high concentration during the experiment. Data on the number of visits per hour per flower were used in a general linear mixed model with Poisson distribution and an observation level random factor to account for overdispersion. The model gives a significant effect of experimental day (*p* = 0.00084), as well as a significant interaction between the Biogluc concentration and experimental day (*p* < 2.2e^-16^) on the number of visits. A significant effect was also found for the circadian rhythm in foraging (*p* < 2.2e^-16^). Post hoc testing (see post hoc table S4) shows an increasing trend (estimate = 0.34, *p* < 0.001) in visitation of the higher versus lower concentration flowers, with a preference tipping point on day three. The effect plot (Fig. 3D) gives a clear view on this preference shift, including the circadian rhythm in foraging. Additional replicate trials are necessary to get more robust results, however, the current results are purely intended as a demonstration of the capabilities of the robotic flower system.

## Discussion

The combination of many automated flowers in an experimental setup creates a robotic flower field, in which every flower is wireless and completely independent. This implies that the setup can easily be changed, even during an experiment: robotic flowers can be moved, added, or removed, and the rate of refilling nectar can be altered remotely. This can be useful in research questions dealing with ephemeral food sources or changing patchy availability of resources. As such, it can be used to assess the effect of nectar traits on the behaviour of floral visitors. Not only sugar composition or concentration, but also amino acids, secondary metabolites, and trace elements in nectar can influence pollinator behaviour (Afik et al., 2008; Nepi et al., 2012; Stevenson et al., 2017). Similarly, human induced disturbances (e.g., pesticides or pollution traces (Lämsä et al., 2018)) or microbial inoculation (e.g., by yeasts or bacteria (Alvarez-Pérez et al., 2012; Pozo et al., 2020)) can affect foraging. All these traits can easily be manipulated in experiments using these robotic flowers to assess their influence on pollinators. A concrete example of a setup like this could be a flight cage containing a bumblebee colony and two types of flowers, one with control nectar and one with nectar containing pesticide traces (see supporting picture S5).

We aimed to create a study system as versatile as possible, but still accessible and easy to construct. The firmware was created in such a way that many hardcoded parameters in the firmware can be easily changed in the adjustable ‘variables’ header file. Besides software changes, hardware can be adapted, e.g., extra batteries and sensors could be used. Measuring temperature, humidity, light intensity, or other factors could give useful information. For example, the foraging behaviour of some bees depends on temperature, solar radiation, humidity, and wind speed (Vicens & Bosch, 2000). Another possibility is to use one device holding these extra sensors as a measuring hub to supply centralized data for all surrounding robotic flowers.

The design of the robotic flower can be adapted to fit requirements for specific research topics. Flower characteristics (e.g., flower disk size, shape, or colour) can be easily altered, e.g., to change search time or handling time. This can be used to investigate associative learning behaviour (Menzel, 1993; Werner et al., 2016), opening the possibility to test the learning abilities of bees associated with stimuli that may not be advertised before collecting the reward (e.g., cues that do not affect palatability or odour). Robotic flowers can be constructed to mimic floral displays for investigating visitor selectivity. For example, damaged flower parts can be a sign of smaller nectar reward and possibly invoke avoidance behaviour in pollinators (Goulson et al., 2007). Not only single but also combinations of manipulations are possible, and even dynamic setups in which the system responds to pollinator behaviour or environmental factors by changing parameters could be used. For example, the robotic flower could be programmed to stop offering nectar after a certain number of visits. Furthermore, a range of floral visitors besides bees can be studied, or a comparison between species can be made. In the first proof of concept, it was already shown that non-bee species, specifically syrphid flies, are attracted to the robotic flower as well. But not only insects can be studied: with the right hardware changes also bigger animals such as mammals (e.g., flower visiting rodents) and birds (e.g., hummingbirds) might be targeted with the robotic flower.

Artificial flower systems of any kind also come with constraints (Essenberg, 2015). Although this robotic flower can be used to address a wide range of research questions, it does have limitations and leaves room for improvement. The robotic flower is made of rigid plastic, which does not resemble natural plant tissue in feel and appearance. It currently also lacks UV-patterns that are present in some flowers to guide pollinators to the reward (Hansen et al., 2012).

This will not keep floral visitors away, but it may slightly reduce the salience of the robotic flower (Chittka et al., 1994). Similarly, floral scents could be added to further approximate natural flower systems or to test the influence of plant volatiles on foraging behaviour.

At present it is not yet possible to automatically discriminate between individual floral visitors and the measured foraging pattern only gives a global view and does not supply information on individual choices. Thus, when used in open field, the distinct species identities of visitors are unknown. Taking a picture of the feeding hole at the beginning of a visit could help tackle this issue, with the downside that the pictures are too large to send over LoRaWAN and therefore would need to be stored locally for later analysis. In addition, it increases the price of the system and could decrease battery life. Another possibility is using a combination of information from different sensors such as weight, wingbeat frequency and/or buzz vibration to deduce the visitor identity, maybe by making use of artificial intelligence software, which would require a more powerful (and more expensive) processor.

The measured probing time can serve as a proxy for the nectar volume that is consumed during a visit. Therefore, the maximum visit duration was set at 250 seconds in this prototype, giving visitors enough time to complete nectar depletion. Longer visits have been proven not to be foraging related, but rather consist of grooming, food pump from crop to ventriculus, or rest (Pozo et al., 2020). Regrettably, all these processes are not subjected to an unambiguous act of crossing the infrared sensor and can be detected only by direct observation.

Data communication with LoRaWAN is very well suited for this kind of application, but, like all (wireless) communication technologies, it also has its limitations and concerns (Adelantado et al., 2017). Packages that are lost during transmission cannot be recovered. This has been compensated in the flower design by asking the backend for a receive-confirmation, and if necessary, repeat the message. Using an external memory (which was already provided in the hardware of this prototype) to store data as a local back-up could be another solution in future versions.

Besides nectar, also pollen is collected by pollinators to fulfil their nutritional requirements (Simpson & Neff, 1981, 1983). For example, buzz pollination can be linked to important crop species such as tomatoes (De Luca & Vallejo-Marin, 2013). Therefore, this field of study could make beneficial use of robotic flowers that can present pollen to visitors (see the system made by Switzer et al. (2019)). The function of presenting pollen was not implemented here, but by combining both rewards in the same robotic flower system information on another level might be obtained. An idea could be to add artificial anthers fitted with the necessary electronics to release pollen when buzz vibration is registered.

Although they may not resemble natural systems, in which countless biotic and abiotic factors are at play, by standardizing factors robotic flowers can help researchers to understand fundamental processes (and their relation to each other) that are at play in the complex web that is pollinator behaviour.

## Conclusions

The presented robotic flower advances the state-of-the-art by supplying a full wireless data transmission mechanism to automatically gather foraging behaviour data. The absence of cables or external control units makes it possible to use the robotic flowers in a flexible way, as well as in more remote experimental setups. Moreover, wireless robotic flowers can be easily connected to a network of other wireless devices, allowing for the scaling up of experiments and the collection of data from multiple locations simultaneously. The system is easily customizable by offering many parameters that can be changed, even during a trial. A dashboard website allows the user to monitor all robotic flowers deployed in a setup and change the nectar refill rate remotely for every flower individually. These capabilities make the flower a versatile tool for behavioural studies with pollinators. It has possible applications in pollination ecology, social and associative learning, conservation biology, biocontrol, and ecotoxicology. Moreover, the flower is robust, relatively easy and cheap to construct and offers the appearance of a commercial product, while being still completely customizable.

## Acknowledgements

In memory of Alejandro Arbeláez García, who contributed in the development and consistently brought a sense of enjoyment to the work environment.

This robotic flower would not have been possible without the inspiration and technical ideas of Prof. Jeff Rozenski. We also want to acknowledge Marc Lambaerts and Thomas Pilkington from FabLab KU Leuven for their help with 3D-printing and Kasper Van Acker for his help with the assembly. We thank Ewoud De Moor for his advice on the firmware and we are grateful for the help received by Hanna Schallauer, Jana Offenberger, and Alfredo Benavente Martinez in carrying-out the proof of concept. This research was partly financed by the Research Foundation Flanders (FWO), in the context of project 11L2423N.

## Data accessibility

The firmware, the Node-RED flows, files for 3D printing and PCB design files are available at GitHub: https://github.com/Kamiel-debeuckelaere/One-Robotic-Flower-Field-To-Study-Them-All.git.

## Authors’ Contributions Statement

M.I.P. and H.J. did the conceptualization and proposed the project. K.D. developed the robotic flower system with help of D.J. on the selection of hardware parts and PCB design. K.D. did the software development with substantial input from D.J. and E.S. The assembly and testing of the robotic flowers were done by K.D., D.J. and M.I.P. In the proof of concept, T.W. played an important part with design and data analysis. The writing of the manuscript was led by K.D. and all co-authors critically contributed to the drafts and gave final approval for publication.

## Supporting information

**S1**: screenshots of the different tabs opened in the frontend application (PNG). Top, left to right: ‘Battery’, ‘Alarms’ and ‘Sleep Time’. Bottom, left to right: ‘Refill Gap’ and ‘Files’

**S2**: animation video (mp4) of 3D-assembly showing how distinct parts of the robotic flower fit together

**S3**: file (PDF) with the user manual containing descriptions and figures to show the robotic flower construction and an elaborate guide to setup trials, information about battery life calculation, and troubleshooting

**S4**: table (PNG) with post hoc contrasts of the proof of concept experiment

**S5**: figure (PNG) of two robotic flower field setups with 6 flowers each as an example of a potential experimental setup that mimics different flower patches (blue and yellow)

**S6**: file (Excel) with measurements of the volume of the prototype nectar cup

**S7**: figure (PNG) of electrical schematic

**S8**: file (Excel) with list of all parts that are needed to build one robotic flower (BOM)

**S9**: proof of concept data (CSV)

## References

Adelantado, F., Vilajosana, X., Tuset-Peiro, P., Martinez, B., Melia-Segui, J., & Watteyne, T. (2017). Understanding the Limits of LoRaWAN [Article]. Ieee Communications Magazine, 55(9), 34–40. https://doi.org/10.1109/mcom.2017.1600613

Afik, O., Dag, A., & Shafir, S. (2008). Honeybee, Apis mellifera, round dance is influenced by trace components of floral nectar. Animal Behaviour, 75(2), 371–377. https://doi.org/10.1016/j.anbehav.2007.04.012

Alvarez-Pérez, S., Herrera, C. M., & de Vega, C. (2012). Zooming-in on floral nectar: a first exploration of nectar-associated bacteria in wild plant communities. FEMS Microbiol Ecol, 80(3), 591–602. https://doi.org/10.1111/j.1574-6941.2012.01329.x

Billiet, A., Meeus, I., Van Nieuwerburgh, F., Deforce, D., Wackers, F., & Smagghe, G. (2016). Impact of sugar syrup and pollen diet on the bacterial diversity in the gut of indoor-reared bumblebees (Bombus terrestris). Apidologie, 47(4), 548–560. https://doi.org/10.1007/s13592-015-0399-1

Calisi, R. M., & Bentley, G. E. (2009). Lab and field experiments: are they the same animal? Horm Behav, 56(1), 1–10. https://doi.org/10.1016/j.yhbeh.2009.02.010

Chittka, L., Shmida, A., Troje, N., & Menzel, R. (1994). Ultraviolet as a component of flower reflections, and the colour perception of Hymenoptera. Vision Res, 34(11), 1489–1508. https://doi.org/10.1016/0042-6989(94)90151-1

Cnaani, J., Thomson, J. D., & Papaj, D. R. (2006). Flower Choice and Learning in Foraging Bumblebees: Effects of Variation in Nectar Volume and Concentration. Ethology, 112(3), 278–285. https://doi.org/10.1111/j.1439-0310.2006.01174.x

Crall, J. D., Gravish, N., Mountcastle, A. M., & Combes, S. A. (2015). BEEtag: A Low-Cost, Image-Based Tracking System for the Study of Animal Behavior and Locomotion. Plos One, 10(9). https://doi.org/10.1371/journal.pone.0136487

Dankert, H., Wang, L., Hoopfer, E. D., Anderson, D. J., & Perona, P. (2009). Automated monitoring and analysis of social behavior in Drosophila. Nature Methods, 6(4), 297–303. https://doi.org/10.1038/nmeth.1310

De Luca, P. A., & Vallejo-Marin, M. (2013). What’s the ‘buzz’about? The ecology and evolutionary significance of buzz-pollination. Current Opinion in Plant Biology, 16(4), 429–435. https://doi.org/10.1016/j.pbi.2013.05.002

de Souza, P., Marendy, P., Barbosa, K., Budi, S., Hirsch, P., Nikolic, N., Gunthorpe, T., Pessin, G., & Davie, A. (2018). Low-Cost Electronic Tagging System for Bee Monitoring. Sensors, 18(7), Article 2124. https://doi.org/10.3390/s18072124

Dell, A. I., Bender, J. A., Branson, K., Couzin, I. D., de Polavieja, G. G., Noldus, L. P. J. J., Pérez-Escudero, A., Perona, P., Straw, A. D., Wikelski, M., & Brose, U. (2014). Automated image-based tracking and its application in ecology. Trends in Ecology & Evolution, 29(7), 417–428. https://doi.org/10.1016/j.tree.2014.05.004

Essenberg, C. J. (2015). Flobots: Robotic flowers for bee behaviour experiments. Journal of Pollination Ecology, 15(1), 1. https://doi.org/10.26786/1920-7603(2015)7

Fry, S. N., Bichsel, M., Müller, P., & Robert, D. (2000). Tracking of flying insects using pan-tilt cameras. Journal of Neuroscience Methods, 101(1), 59–67. https://doi.org/10.1016/S0165-0270(00)00253-3

Goulson, D. (2003). Conserving wild bees for crop pollination [Article]. Journal of Food Agriculture & Environment, 1(1), 142–144. https://doi.org/10.1080/00218839.2022.2046528

Goulson, D. (2010). Bumblebees: Behaviour and ecology (2 ed.). Oxford University Press.

Goulson, D., & Cory, J. S. (1993). Flower constancy and learning in foraging preferences of the green-veined white butterfly Pleris napi [10.1111/j.1365-2311.1993.tb01107.x]. Ecological Entomology, 18(4), 315–320. https://doi.org/https://doi.org/10.1111/j.1365-2311.1993.tb01107.x

Goulson, D., Cruise, J. L., Sparrow, K. R., Harris, A. J., Park, K. J., Tinsley, M. C., & Gilburn, A. S. (2007). Choosing rewarding flowers; perceptual limitations and innate preferences influence decision making in bumblebees and honeybees. Behavioral Ecology and Sociobiology, 61(10), 1523–1529. https://doi.org/10.1007/s00265-007-0384-4

Greggers, U., & Menzel, R. (1993). Memory Dynamics and Foraging Strategies of Honeybees [Article]. Behavioral Ecology and Sociobiology, 32(1), 17–29. https://doi.org/10.1007/BF00172219

Gumbert, A. (2000). Color choices by bumble bees (Bombus terrestris): innate preferences and generalization after learning. Behavioral Ecology and Sociobiology, 48(1), 36–43. https://doi.org/10.1007/s002650000213

Hagen, M., Wikelski, M., & Kissling, W. D. (2011). Space Use of Bumblebees (Bombus spp.) Revealed by Radio-Tracking. Plos One, 6(5), Article e19997. https://doi.org/10.1371/journal.pone.0019997

Hammer, M., & Menzel, R. (1995). Learning and memory in the honeybee. The Journal of Neuroscience, 15(3), 1617–1630. https://doi.org/10.1523/jneurosci.15-03-01617.1995

Hannah, L., Dyer, A. G., Garcia, J. E., Dorin, A., & Burd, M. (2019). Psychophysics of the hoverfly: categorical or continuous color discrimination? Current Zoology, 65(4), 483–492. https://doi.org/10.1093/cz/zoz008

Hansen, D. M., Van der Niet, T., & Johnson, S. D. (2012). Floral signposts: testing the significance of visual ‘nectar guides’ for pollinator behaviour and plant fitness. Proc Biol Sci, 279(1729), 634–639. https://doi.org/10.1098/rspb.2011.1349

Heinrich, B. (1981). Bumblebee economics (3 ed.). Harvard University Press.

Ingram, E. M., Augustin, J., Ellis, M. D., & Siegfried, B. D. (2015). Evaluating sub-lethal effects of orchard-applied pyrethroids using video-tracking software to quantify honey bee behaviors. Chemosphere, 135, 272–277. https://doi.org/10.1016/j.chemosphere.2015.04.022

Ings, T. C., & Chittka, L. (2008). Speed-Accuracy Tradeoffs and False Alarms in Bee Responses to Cryptic Predators. Current Biology, 18(19), 1520–1524. https://doi.org/10.1016/j.cub.2008.07.074

Keasar, T. (2000). The spatial distribution of nonrewarding artificial flowers affects pollinator attraction [Article]. Animal Behaviour, 60, 639–646. https://doi.org/10.1006/anbe.2000.1484

Klein, A. M., Vaissiere, B. E., Cane, J. H., Steffan-Dewenter, I., Cunningham, S. A., Kremen, C., & Tscharntke, T. (2007). Importance of pollinators in changing landscapes for world crops [Review]. Proceedings of the Royal Society B-Biological Sciences, 274(1608), 303–313. https://doi.org/10.1098/rspb.2006.3721

Kuusela, E., & Lämsä, J. (2016). A low-cost, computer-controlled robotic flower system for behavioral experiments [Article]. Ecology and Evolution, 6(8), 2594–2600. https://doi.org/10.1002/ece3.2062

Lämsä, J., Kuusela, E., Tuomi, J., Juntunen, S., & Watts, P. C. (2018). Low dose of neonicotinoid insecticide reduces foraging motivation of bumblebees. Proceedings of the Royal Society B-Biological Sciences, 285(1883), Article 20180506. https://doi.org/10.1098/rspb.2018.0506

Lunau, K. (2000). The ecology and evolution of visual pollen signals. Plant Systematics and Evolution, 222(1), 89–111. https://doi.org/10.1007/BF00984097

Lunau, K., Wacht, S., & Chittka, L. (1996). Colour choices of naive bumble bees and their implications for colour perception. Journal of Comparative Physiology a-Neuroethology Sensory Neural and Behavioral Physiology, 178(4), 477–489. https://doi.org/10.1007/BF00190178

Maggiora, R., Saccani, M., Milanesio, D., & Porporato, M. (2019). An Innovative Harmonic Radar to Track Flying Insects: the Case of Vespa velutina. Scientific Reports, 9(1), 11964. https://doi.org/10.1038/s41598-019-48511-8

Makino, T. T., & Sakai, S. (2007). Experience changes pollinator responses to floral display size: from size-based to reward-based foraging [Article]. Functional Ecology, 21(5), 854–863. https://doi.org/10.1111/j.1365-2435.2007.01293.x

Makinson, J. C., Woodgate, J. L., Reynolds, A., Capaldi, E. A., Perry, C. J., & Chittka, L. (2019). Harmonic radar tracking reveals random dispersal pattern of bumblebee (Bombus terrestris) queens after hibernation. Scientific Reports, 9, Article 4651. https://doi.org/10.1038/s41598-019-40355-6

Menzel, R. (1993). Associative learning in honey bees. Apidologie, 24(3), 157–168. https://doi.org/10.1051/apido:19930301

Minahan, D. F., & Brunet, J. (2018). Strong Interspecific Differences in Foraging Activity Observed Between Honey Bees and Bumble Bees Using Miniaturized Radio Frequency Identification (RFID). Frontiers in Ecology and Evolution, 6, Article 156. https://doi.org/10.3389/fevo.2018.00156

Moffatt, L. (2001). Metabolic rate and thermal stability during honeybee foraging at different reward rates. J Exp Biol, 204(Pt 4), 759–766. https://doi.org/10.1242/jeb.204.4.759

Nachev, V., & Winter, Y. (2012). The psychophysics of uneconomical choice: non-linear reward evaluation by a nectar feeder [Article]. Animal Cognition, 15(3), 393–400. https://doi.org/10.1007/s10071-011-0465-7

Nepi, M., Soligo, C., Nocentini, D., Abate, M., Guarnieri, M., Cai, G., Bini, L., Puglia, M., Bianchi, L., & Pacini, E. (2012). Amino acids and protein profile in floral nectar: Much more than a simple reward. Flora - Morphology, Distribution, Functional Ecology of Plants, 207(7), 475–481. https://doi.org/10.1016/j.flora.2012.06.002

Nicolson, S. W. (2011). Bee Food: The Chemistry and Nutritional Value of Nectar, Pollen and Mixtures of the Two. African Zoology, 46(2), 197–204, 198. https://doi.org/10.3377/004.046.0201

Nicolson, S. W., & Thornburg, R. W. (2007). Nectar chemistry. In S. W. Nicolson, M. Nepi, & E. Pacini (Eds.), Nectaries and Nectar (pp. 215–264). Springer Netherlands. https://doi.org/10.1007/978-1-4020-5937-7_5

Nunes-Silva, P., Hrncir, M., Guimaraes, J. T. F., Arruda, H., Costa, L., Pessin, G., Siqueira, J. O., de Souza, P., & Imperatriz-Fonseca, V. L. (2019). Applications of RFID technology on the study of bees. Insectes Sociaux, 66(1), 15–24. https://doi.org/10.1007/s00040-018-0660-5

Ohashi, K., D’Souza, D., & Thomson, J. D. (2010). An automated system for tracking and identifying individual nectar foragers at multiple feeders. Behavioral Ecology and Sociobiology, 64(5), 891–897. https://doi.org/10.1007/s00265-010-0907-2

Ollerton, J., Winfree, R., & Tarrant, S. (2011). How many flowering plants are pollinated by animals? Oikos, 120(3), 321–326. https://doi.org/10.1111/j.1600-0706.2010.18644.x

Paldi, N., Zilber, S., & Shafir, S. (2003). Associative olfactory learning of honeybees to differential rewards in multiple contexts--effect of odor component and mixture similarity. J Chem Ecol, 29(11), 2515–2538. https://doi.org/10.1023/a:1026362018796

Potts, S. G., Biesmeijer, J. C., Kremen, C., Neumann, P., Schweiger, O., & Kunin, W. E. (2010). Global pollinator declines: trends, impacts and drivers. Trends in Ecology & Evolution, 25(6), 345–353. https://doi.org/10.1016/j.tree.2010.01.007

Pozo, M. I., van Kemenade, G., van Oystaeyen, A., Aledón-Catalá, T., Benavente, A., Van den Ende, W., Wäckers, F., & Jacquemyn, H. (2020). The impact of yeast presence in nectar on bumble bee behavior and fitness. Ecological Monographs, 90(1), 1–22. https://doi.org/10.1002/ecm.1393

Provoost, M., & Weyns, D. (2019, 25-25 May 2019). DingNet: A Self-Adaptive Internet-of-Things Exemplar. 2019 IEEE/ACM 14th International Symposium on Software Engineering for Adaptive and Self-Managing Systems (SEAMS),

R Core Team. (2022). R: A language and environment for statistical computing. In R Foundation for Statistical Computing. https://www.R-project.org/

Ratnayake, M. N., Dyer, A. G., & Dorin, A. (2021). Tracking individual honeybees among wildflower clusters with computer vision-facilitated pollinator monitoring. Plos One, 16(2). https://doi.org/10.1371/journal.pone.0239504

Real, L., & Rathcke, B. J. (1988). Patterns of Individual Variability in Floral Resources. Ecology, 69(3), 728–735. https://doi.org/10.2307/1941021

Real, L. A. (1981). Uncertainty and Pollinator-Plant Interactions: The Foraging Behavior of Bees and Wasps on Artificial Flowers. Ecology, 62(1), 20–26. https://doi.org/10.2307/1936663

Riley, J. R., Smith, A. D., Reynolds, D. R., Edwards, A. S., Osborne, J. L., Williams, I. H., Carreck, N. L., & Poppy, G. M. (1996). Tracking bees with harmonic radar. Nature, 379(6560), 29–30. https://doi.org/10.1038/379029b0

Schmid-hempel, P., & Stauffer, H. (1998). Parasites and flower choice of bumblebees. Anim Behav, 55(4), 819–825. https://doi.org/10.1006/anbe.1997.0661

Shearwood, J., Aldabashi, N., Eltokhy, A., Franklin, E. L., Raine, N. E., Zhang, C., Palmer, E., Cross, P., & Palego, C. (2021). C-Band Telemetry of Insect Pollinators Using a Miniature Transmitter and a Self-Piloted Drone. IEEE Transactions on Microwave Theory and Techniques, 69(1), 938–946. https://doi.org/10.1109/TMTT.2020.3034323

Simons, D. J., & Chabris, C. F. (1999). Gorillas in Our Midst: Sustained Inattentional Blindness for Dynamic Events. Perception, 28(9), 1059–1074. https://doi.org/10.1068/p281059

Simpson, B. B., & Neff, J. L. (1981). Floral Rewards: Alternatives to Pollen and Nectar. Annals of the Missouri Botanical Garden, 68(2), 301–322. https://doi.org/10.2307/2398800

Simpson, B. B., & Neff, J. L. (1983). Floral Biology and Floral Rewards of Lysimachia (Primulaceae). The American Midland Naturalist, 110(2), 249–256. https://doi.org/10.2307/2425266

Sokolowski, M. B. C., & Abramson, C. I. (2010). From foraging to operant conditioning: A new computer-controlled Skinner box to study free-flying nectar gathering behavior in bees. Journal of Neuroscience Methods, 188(2), 235–242. https://doi.org/10.1016/j.jneumeth.2010.02.013

Spaethe, J., Tautz, J., & Chittka, L. (2001). Visual constraints in foraging bumblebees: Flower size and color affect search time and flight behavior. Proceedings of the National Academy of Sciences of the United States of America, 98(7), 3898–3903. https://doi.org/10.1073/pnas.071053098

Spaethe, J., & Weidenmüller, A. (2002). Size variation and foraging rate in bumblebees (Bombus terrestris). Insectes Sociaux, 49(2), 142–146. https://doi.org/10.1007/s00040-002-8293-z

Stevenson, P. C., Nicolson, S. W., & Wright, G. A. (2017). Plant secondary metabolites in nectar: impacts on pollinators and ecological functions. Functional Ecology, 31(1), 65–75. https://doi.org/10.1111/1365-2435.12761

Sun, S., Leshowitz, M. I., & Rychtář, J. (2018). The signalling game between plants and pollinators. Scientific Reports, 8(1), 6686. https://doi.org/10.1038/s41598-018-24779-0

Switzer, C. M., Russell, A. L., Papaj, D. R., Combes, S. A., & Hopkins, R. (2019). Sonicating bees demonstrate flexible pollen extraction without instrumental learning [Article]. Current Zoology, 65(4), 425–436. https://doi.org/10.1093/cz/zoz013

Thomson, J. D., Ogilvie, J. E., Makino, T. T., Arisz, A., Raju, S., Rojas-Luengas, V., & Tan, M. (2012). Estimating pollination success with novel artificial flowers: Effects of nectar concentration. Journal of Pollination Ecology, 9(0), 108–114. https://doi.org/10.26786/1920-7603(2012)14

Van Geystelen, A., Benaets, K., de Graaf, D. C., Larmuseau, M. H. D., & Wenseleers, T. (2016). Track-a-Forager: a program for the automated analysis of RFID tracking data to reconstruct foraging behaviour. Insectes Sociaux, 63(1), 175–183. https://doi.org/10.1007/s00040-015-0453-z

Vicens, N., & Bosch, J. (2000). Weather-Dependent Pollinator Activity in an Apple Orchard, with Special Reference to Osmia cornuta and Apis mellifera (Hymenoptera: Megachilidae and Apidae). Environmental Entomology, 29(3), 413–420. https://doi.org/10.1603/0046-225x-29.3.413

Waddington, K. D., & Holden, L. R. (1979). Optimal Foraging - Flower Selection by Bees [Article]. American Naturalist, 114(2), 179–196. https://doi.org/10.1086/283467

Werner, A., Stürzl, W., & Zanker, J. (2016). Object Recognition in Flight: How Do Bees Distinguish between 3D Shapes? Plos One, 11(2). https://doi.org/10.1371/journal.pone.0147106

Woodgate, J. L., Makinson, J. C., Lim, K. S., Reynolds, A. M., & Chittka, L. (2016). Life-Long Radar Tracking of Bumblebees. Plos One, 11(8), Article e0160333. https://doi.org/10.1371/journal.pone.0160333

