## Supplementary figures and images for "A wireless, user-friendly, and unattended robotic flower system to assess pollinator foraging behaviour"

### Supplemental figure 1

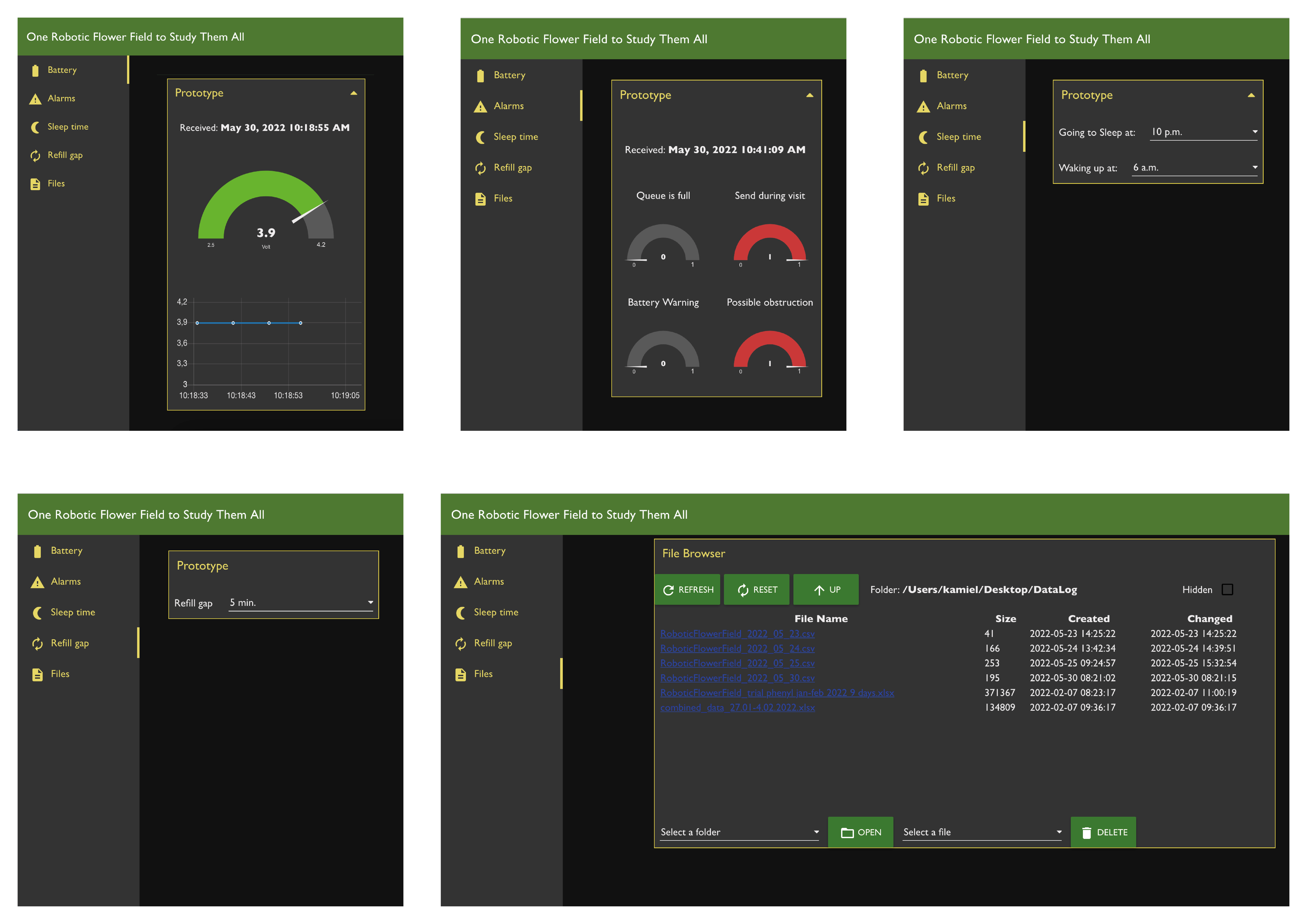

### Supplemental figure 5

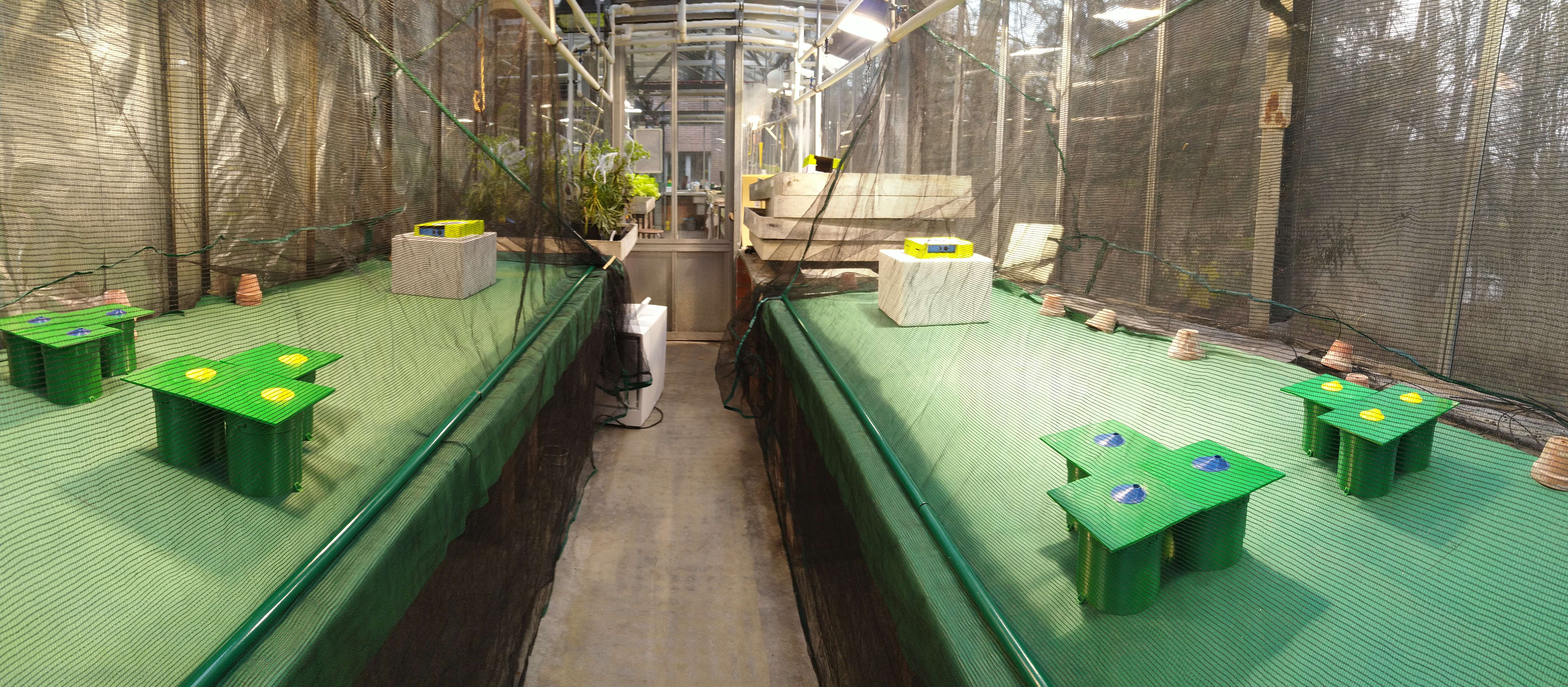

### Supplemental figure 7

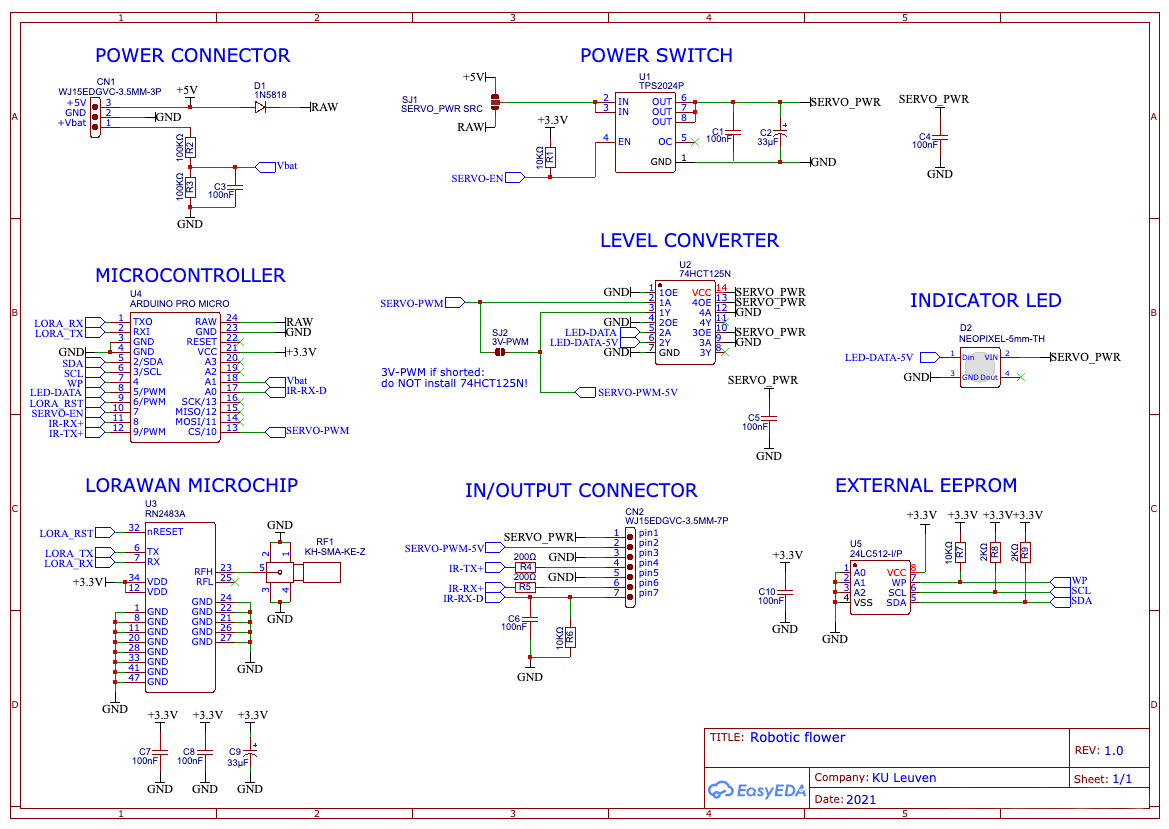

### Supplemental table 4

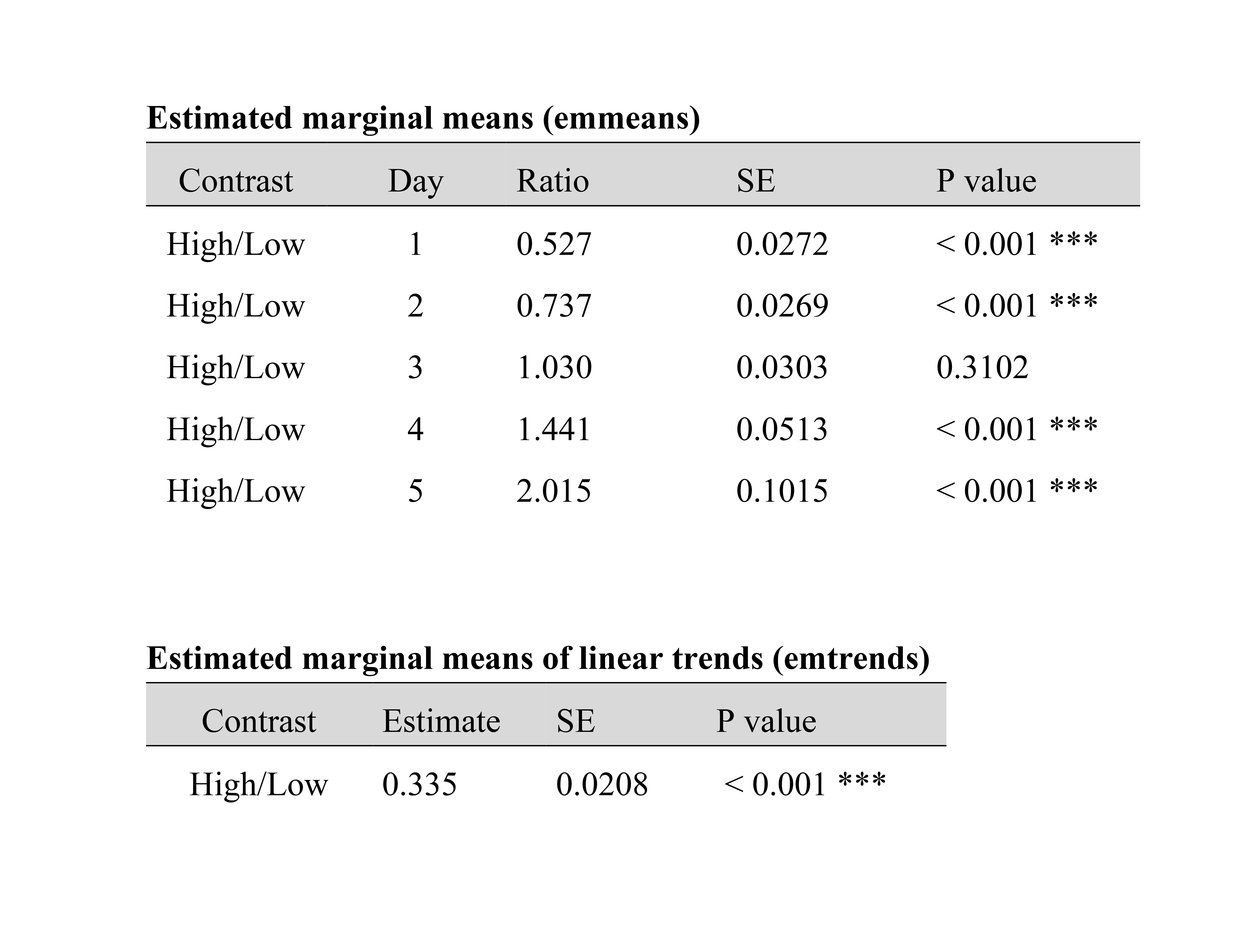
