## Supplemental file 3 for "A wireless, user-friendly, and unattended robotic flower system to assess pollinator foraging behaviour"

### One Robotic Flower Field to Study Them All

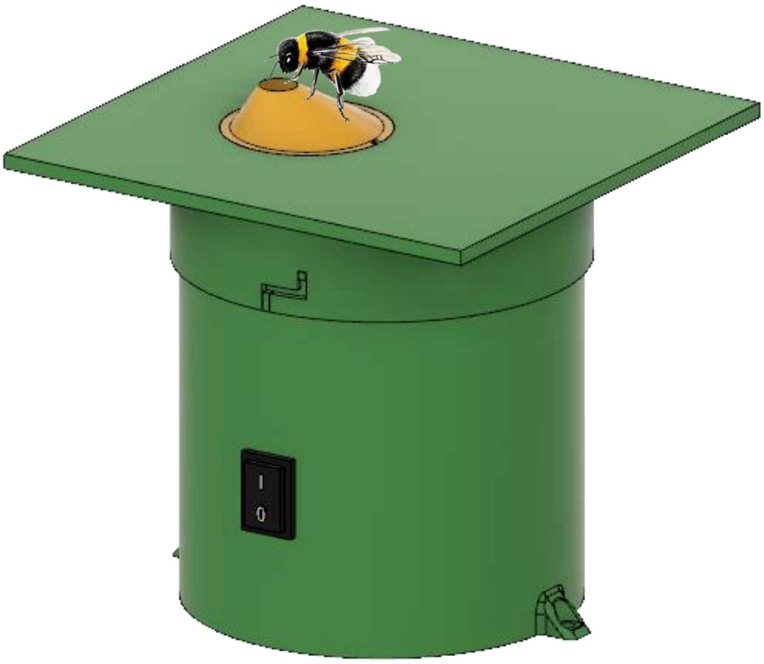

#### User manual

### 1 INTRODUCTION

Assembling the flower(s), creating your own Node-Red dashboard, uploading the firmware to the Arduino… It sounds like a lot of work, but by making this guide we aim to help you to get this done before you know it. You will have to make sure everything works as it is supposed to before the final step: preparing the experimental set-up. We will try to lead you through this process and give some tips that can save some trouble when using this robotic flower system. And in the end, our only hope is that the robotic flowers will prove very helpful in gathering the data that you want!

As a first step, you have to go to the GitHub repository: https://github.com/Kamiel-debeuckelaere/One-Robotic-Flower-Field-To-Study-Them-All . In there, you can download a zip-folder containing all documents to build this robotic flower by clicking on the green button saying 'Code' (Fig. 0). Keep the structure with subfolders and save it to an accessible part of your computer (so that it is easy to access afterwards). These subfolders will be used for different parts in this user manual.

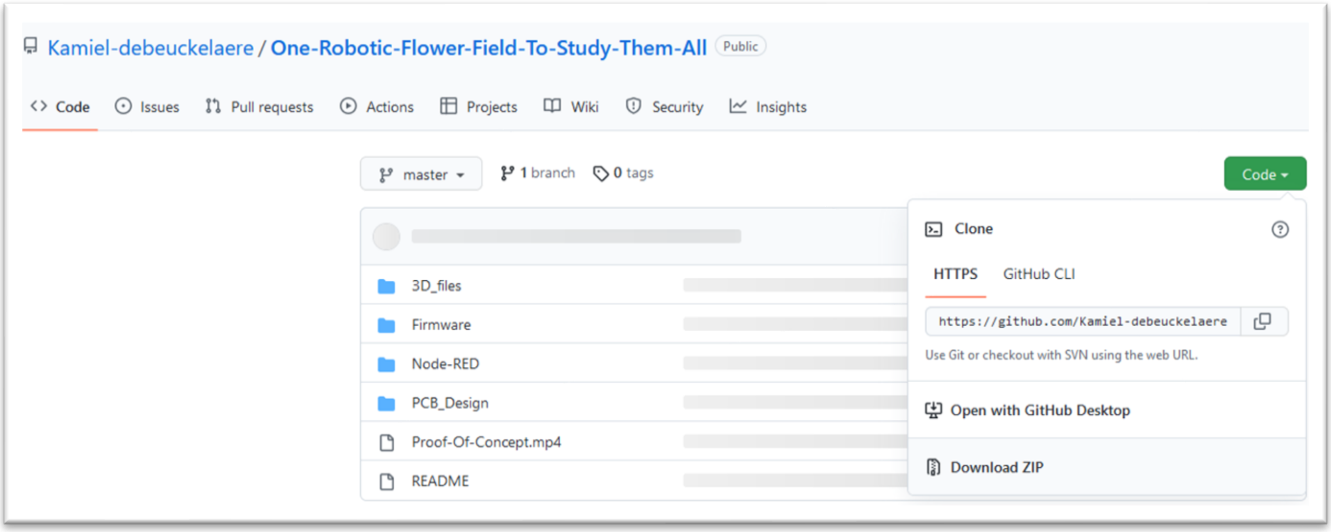

**Figure 0**

### 2 ROBOTIC FLOWER CONSTRUCTION

Make sure you have all materials that are needed to construct the robotic flowers. In the parts list (appendix 1) you can find every item that is needed. You can also find the store and the link to the product we have used, but alternatives are possible. Also make sure you have all the necessary tools to complete the construction, or at least have access to them (e.g., soldering station, 3D-printer, electrical screwdriver etc.).

#### 2.1 PCB

- To print the PCB, you will need the Gerber files of the design. To get these, the .json files on GitHub must be opened with the online (and free) software EasyEDA, after creating an account. The PCB's can be ordered directly through this website as well.

- EasyEDA: https://easyeda.com/

- When PCB's have arrived, carefully solder all the needed parts to the PCB. To do this, see electrical scheme (appendix 2). We started with the through hole parts (ordered from small to large parts for practical reasons during soldering) and ended with the surface mounted LoRa module (rn2483a). A useful tip might be to use a stereomicroscope to solder the relatively fine connections of this LoRa module, as any wrong connection will lead to malfunction. The soldering of the LoRa module is a precision job and can be a bit of a challenge, but it is possible!

- For normal use with battery, connect the bottom two pads of solder jumper SJ1 with tin. In this way, power from the battery is directed to the power switch. When you want to use the flowers connected to the power grid with power coming from the micro-USB on the microcontroller, the top 2 pads must be connected with tin.

- Here, we use a 5V PWM servomotor, meaning solder jumper SJ2 can be ignored. When working with a 3.3V PWM servomotor, SJ2 needs to be connected.

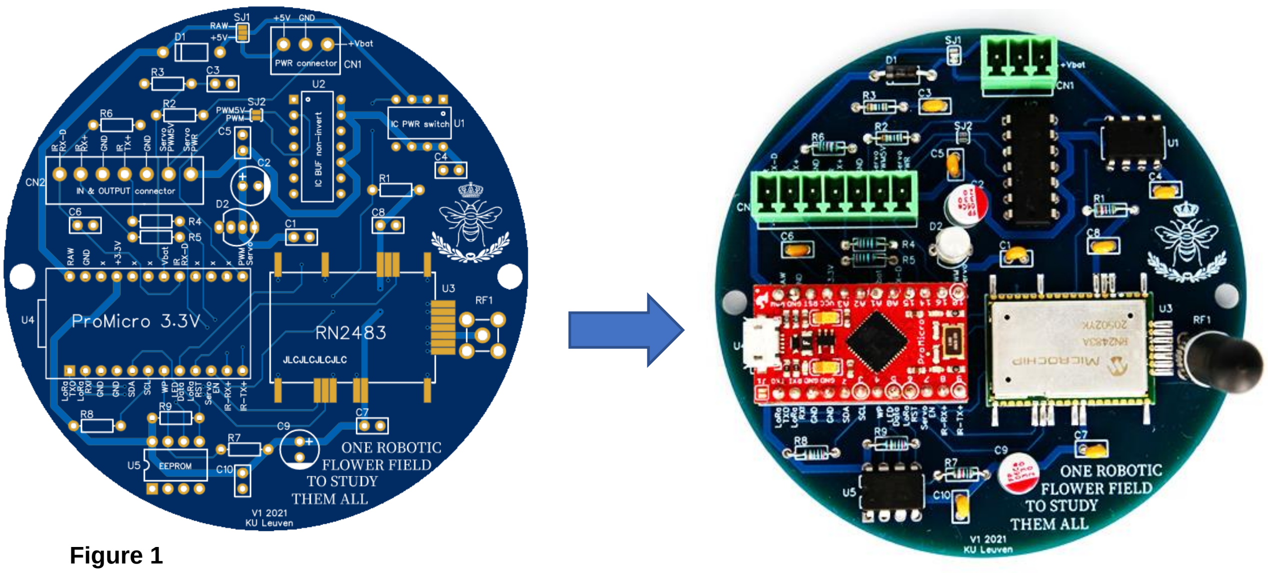

**Figure 1**

#### 2.2 BATTERY MODULE

- Solder the connector to the battery module board and plug in the cables going to the rocker switch. The rocker switch itself can be connected later (see Fig. 3).

- Attach a wire (we used 25 cm) to one of the cables (cut and solder together) going to the rocker switch. This is to let the Arduino read the battery voltage. To cover the part of the cable where now the insulation (plastic cover) is missing, you can use heat-shrink tubing (see Fig. 2).

- Solder wires (positive + and negative -) to the UPS (uninterrupted power source) output of the board. Make sure these wires are long enough (we used 20 cm).

- Solder the board to the battery holder with through-hole soldering and make sure the + and – side are correct

- The clip-in part of the module was originally designed for flat-top batteries but was here manually modified with pincers to fit the button-top battery we selected. If used with flat-top batteries, no modification would be needed.

- Connect the +, - and voltage reader at the correct place to the power connector (term block) using a small screwdriver. This way, the wires can be plugged into to the PCB later.

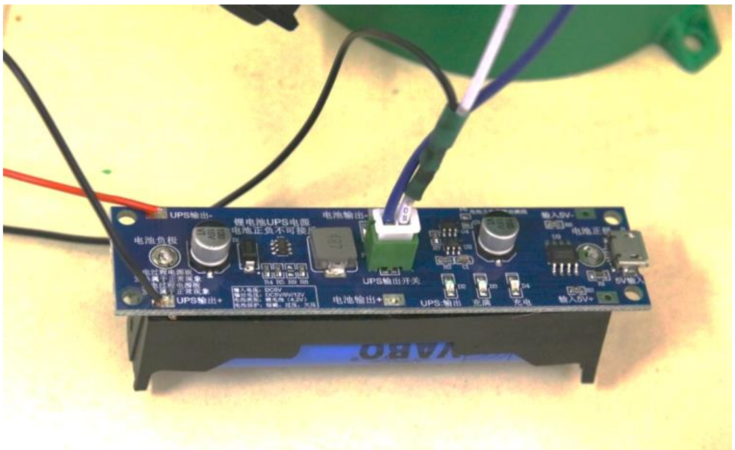

**Figure 2**

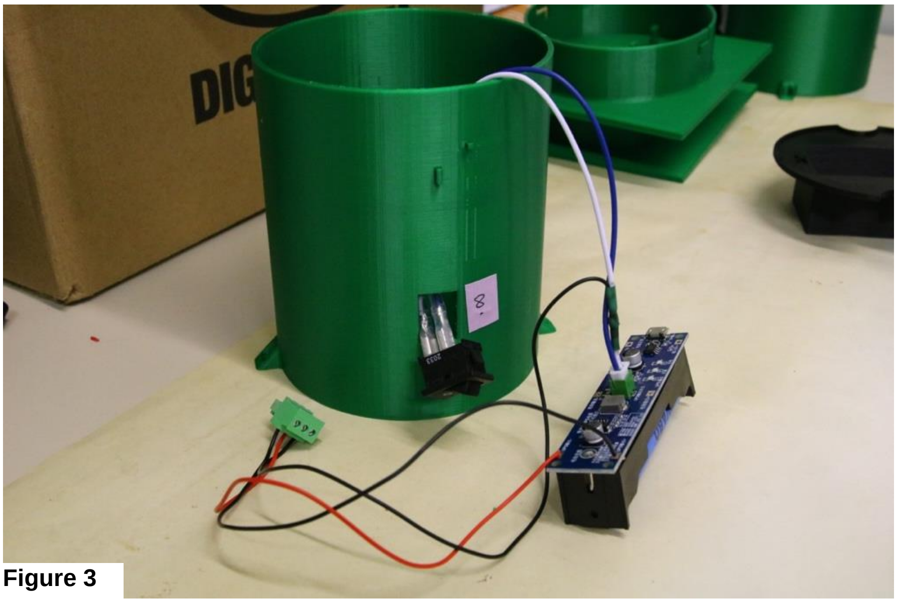

**Figure 3**

#### 2.3 3D-PRINTING & ASSEMBLY

The STL files for 3D printing can be found in GitHub (https://github.com/Kamiel-debeuckelaere/One-Robotic-Flower-Field-To-Study-Them-All/tree/master/3D_files). If you wish to modify parts, you can download and install Autodesk Fusion 360 (https://www.autodesk.com/products/fusion-360/overview?term=1-YEAR&tab=subscription), a 3D designing software which has a free hobby version and a free educational version (for qualifying students and teachers). In this program, you can then upload the zip compressed archive file (.f3z) and start editing. Moreover, many tutorials on how to use this software can be found online.

The robotic flower has 5 3D-printed parts: central flower disk (Fig. 4, A), nectar cup (Fig. 4, B), background plane (Fig. 4, C), reservoir case (Fig. 4, D) and the stem (Fig. 4, E). After printing, first carefully remove the support structures (the most parts can be done manually, but for smaller edges e.g., by using a needle nose plier or a scalpel), and make sure there is no hindering for assembly anymore. Dimensions of the robotic flower can be seen in Fig. 5. We recommend watching also the short assembly animation movie that can be found on GitHub.

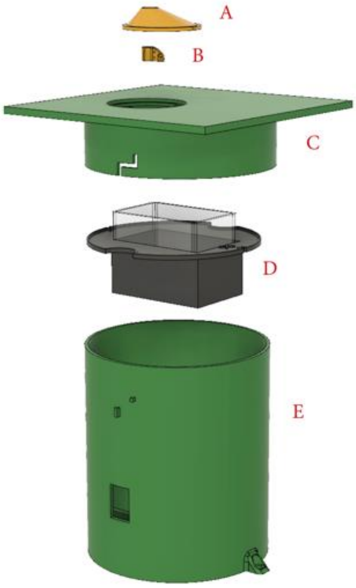

**Figure 4**

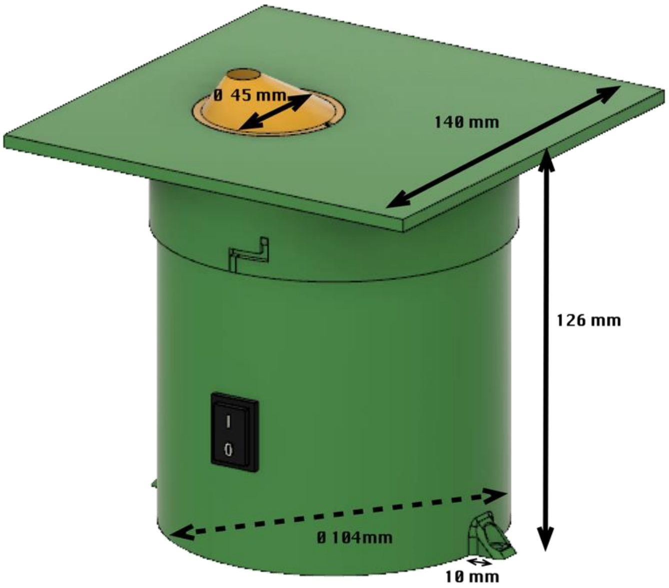

**Figure 5**

##### 2.3.1 STEM

- At the bottom, four beams (Fig. 6, B) are in place to support the PCB so that it is elevated from the bottom. This was done to prevent the PCB being in contact with water when condensation is formed in the flower and runs down the sides. A slit between the wall and PCB is kept free so that the condensation can run to the bottom. To keep the PCB in place, two of the four supports have cylindrical protrusions that fit the holes of the PCB.

- On top of the PCB, there is space left to place the battery module with battery. This design with a broad stem, in which the battery is not put upright, ensures a low centre of gravity, helping to avoid easy tipping over of the robotic flower.

- The stem has two support beams (Fig. 6, A), one at each side to hold up the nectar reservoir case (see further). Two protrusions are provided which fit the notches in the reservoir case to prevent it from moving around. Keeping the right position is important for the well-functioning of the servo refilling system.

- In the side of the stem there is a hole (Fig. 6, C) to fit in the on/off switch that can be clipped in.

- Protrusions for the twist-and-lock system (Fig. 6, D) to fit the background plane on top of the stem which keeps the components from disconnecting easily.

- The mounting feet (Fig. 6, E) are constructed to fix the robotic flower on the substrate to keep it from tipping over if necessary. For example, the flower could be anchored in the ground using a small picket, or it could be screwed onto a platform.

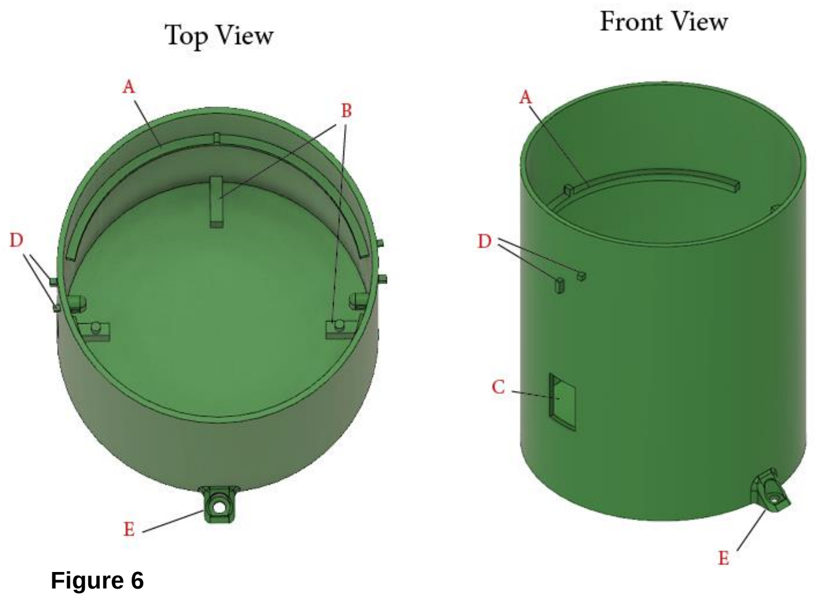

**Figure 6**

##### 2.3.2 BACKGROUND PLANE AND CENTRAL FLOWER DISK

- The background plane is constructed so that it fits over the stem and is kept in place with a twist-and-lock system for which notches are present on the side (Fig. 7, A).

- On the bottom of the background plane two protrusions with holes are constructed (Fig. 7, B) to attach the servomotor (Fig. 7, C) with zip ties (2.5 x 200 mm).

- In the place where the central flower disk must fit, an opening with a ledge is made. In this ledge, a protrusion is constructed in such a way that the central flower disk (Fig. 7, D) with notch can be placed in the right position before being fixated with transparent silicone. In this way, the seal is also made waterproof.

- In the central flower disk, a cylindrical hole is constructed through which pollinators can enter the flower to reach the nectar. The feeding hole can thus not be too small for the large workers to be able to enter it. Still, in contrast to the 13 mm in the previous version of Kuusela & Lämsä (2016), we chose a smaller diameter for the feeding hole in this new robotic flower. Based on experience by Pozo et al. (2020) that showed a mean *Bombus terrestris* worker thorax width of 6.97 ± 0.85 mm we chose to make the feeding hole 10 mm in diameter to reduce the interference of external light (e.g., from the sun) with the infrared sensor while still leaving enough space for the workers to enter.

- The depth of the feeding hole should be deep enough so that visitors are forced to crawl inside to reach the nectar and thereby causing a disturbance in the infrared barrier of the detection system. With the average length of *B. terrestris* workers being 14.1 ± 2.4 mm (personal communication with Biobest, Westerlo, Belgium), the depth of the feeding hole was chosen to be 9 mm.

- The infrared (IR) emitter and sensor both have a positive and negative pole, which are soldered with one wire each (black for – and red for +; we used a wire-length of 40 mm) and covered with heat-shrink tubing. The IR emitter and sensor are then glued in the two designated holes in the central flower disk (Fig. 7, E; Fig. 8) with transparent silicone. The 4 wires coming from the from the IR emitter and sensor are kept together with heat-shrink tubing and are attached to the term block together with the 3 wires coming from the servomotor.

- At the bottom of the feeding hole (Fig. 7, F), there is a nectar hole with a diameter of 3 mm and depth of 1 mm through which floral visitors can reach the nectar cup that is pressed against the central flower disk by the servomotor. A notch shaped like the nectar cup is made at the bottom of the central flower disk to ensure a tight fit and a correct position of the nectar cup relative to the central flower disk.

- The conical shape of the central flower disk is to avoid rain running off from the background plane in the feeding hole as much as possible, but the shape surrounding the feeding hole can be changed.

- Other structures are protrusions with holes to guide the cables coming from the infrared emitter and sensor so that they don't hinder the servo-arm during refilling (Fig. 7, G).

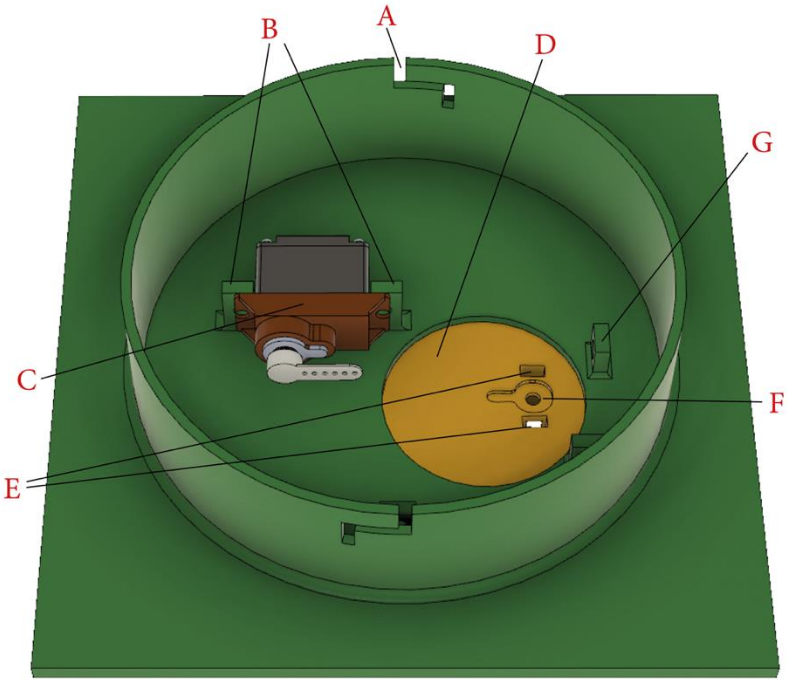

**Figure 7**

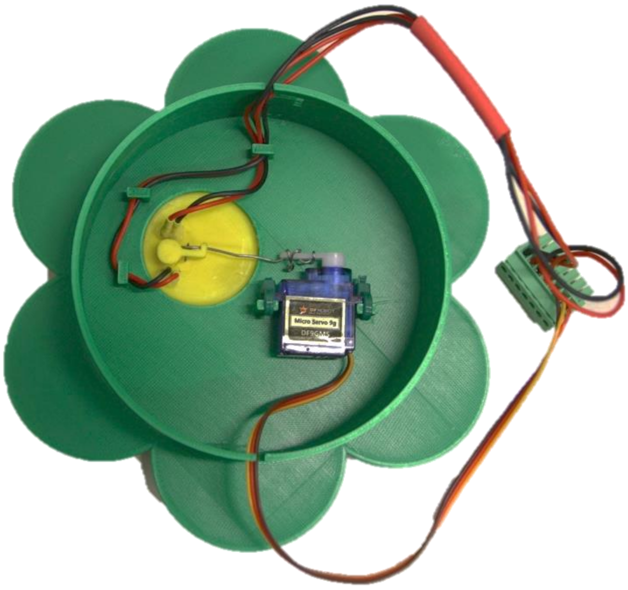

**Figure 8**

##### 2.3.3 NECTAR CUP

- By changing the 3D-design of the nectar cup, it is easy to adapt the discrete volume of the nectar presented to visitors of the robotic flower according to specific needs. For this prototype version, the volume is approximately 15µL (see repeated measurements with capillary tubes in supporting information of the paper).

- Another characteristic to consider here is the length of the proboscis of visitors, this because they need to be able to reach the bottom part of the nectar cup. The proboscis length varies among species and is an ecological important factor in assessing which species have access to floral rewards. The hole in this nectar cup (Fig. 9, A) has a diameter of 3 mm (to match the hole in the central flower disk) and depth of 3.5 mm. The thickness of the central flower disk at the point where it meets the nectar cup is 1 mm. This distance must be added to the depth of the hole in the nectar cup, making a total depth of 4.5 mm. This is easily reachable for *B. terrestris* (and *B. lucorum*) workers with a functional proboscis length of 8.1 ± 0.2 mm, *Apis mellifera* workers with 6.6 ± 032 mm and *B. pascuorum* workers with 8.9 ± 0.2 mm (Balfour et al., 2013). The proboscis length of Lepidoptera in a study with 15 species by Corbet (2000) also showed to be sufficient to reach the nectar in the robotic flower, ranging from 7.34 ± 0.07 mm for *Lycaena phlaeas* to 16.19 ± 0.41 mm for *Inachis io*. Most Syrphinae (Diptera) have a proboscis that is relatively short related to their size, but some species, including *Eupeodes corollae* and *Platycheirus manicatus* have a longer proboscis (3-5 mm) (Branquart & Hemptinne, 2000). For these syrphids, the nectar cup might be too deep to reach all the way to the bottom, but during the proof of concept this did not seem to stop them from visiting the robotic flower.

- The nectar cup has three holes (Fig. 9, B) through which the part of the servo-arm, consisting of steel wire (diameter 0.8 mm), can be attached. The other end of the servo-arm is clipped on the spindle of the servomotor. Because the servo-arm partly consists of steel wire, it can be manipulated manually to achieve a close fit between the nectar cup and the central flower disk, so that both holes are matching (see Fig. 8). More information in part '7.2 Attaching servo arm'.

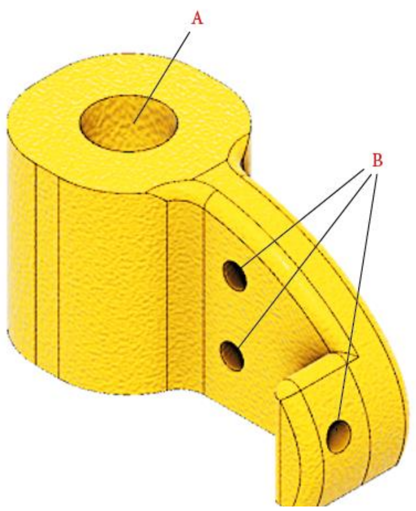

**Figure 9**

##### 2.3.4 Reservoir case

- The reservoir case is built to carry the nectar reservoir (Fig. 10, E) while also forming a protective barrier between the electronic components at the bottom and the artificial nectar located at the top.

- The case is supported by beams on the stem and is kept in the right position by notches (Fig. 10, B) that fit in protrusions of these beams.

- Crescent notches (Fig. 10, C) at both sides of the nectar reservoir case ensure a passage for the cables from the servomotor, infrared sensor and emitter attached to the bottom of the background plane running towards the electronics compartment at the bottom of the stem.

- A rim (Fig. 10, D) is provided on the edge of the reservoir case to prevent spill-over nectar from the nectar reservoir to seep into the electronics compartment underneath. In the latest updated version of the 3D-design, these edges have been made higher than is seen in Fig. 10.

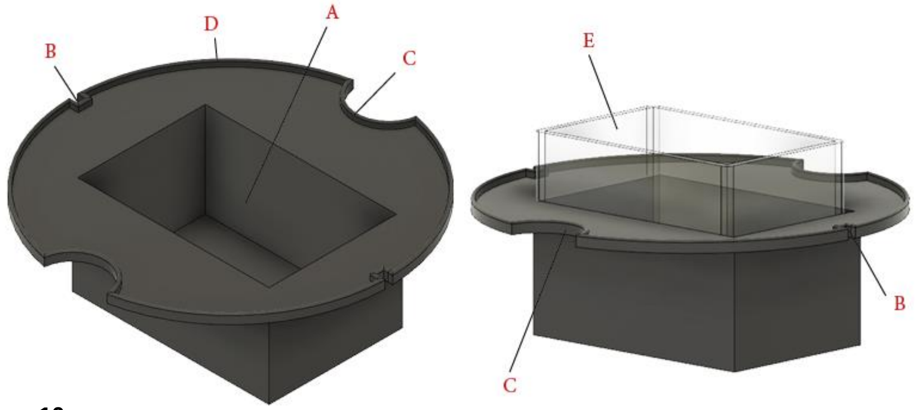

**Figure 10**

##### 2.3.5 NECTAR RESERVOIR

- For this prototype an off-the-shelf reservoir with inside dimensions of 56 × 36 × 44 mm (L × B × H) was used, giving it a volume of 88.7 ml (Fig. 10, E).

- In a hypothetical set-up with a nectar cup of 25 µL that is depleted every time, a standard refilling rate of 10 minutes and in working mode for 14 hours every day, the robotic flower would theoretically be capable of offering nectar for more than 40 days. This is when the nectar reservoir is filled up completely to the edge, ignoring spill-over and evaporation and assuming the servo-refilling system can scoop all the nectar down to the last drop. This is not a realistic scenario, but it shows that there is enough margin for an experimental set-up of e.g., two weeks. (With long experimental periods other problems will probably arise, such as uncontrolled bacterial growth in the artificial nectar.)

- If another size is wanted, a nectar container could also be custom made, with the restriction that also the nectar reservoir case will have to be modified and keeping in mind that the servo-arm must fit in the reservoir when refilling.

### 3 SETUP IOT

To work with Internet of Things, you will need a LoRaWAN backend and one ore multiple gateways, depending on the specific application. If you are lucky, you might use a pre-existing LoRaWAN communication infrastructure provided by your university or institute (as we did). We recommend you to inquire about this at your ICT service.

If no such service is available to you (yet), it is still possible to get a gateway yourself and use 'The Things Network', which provides a free LoRaWAN backend infrastructure globally. More information can be found here: https://www.thethingsnetwork.org/

You will have to make an application (choose unique name) first. Here you can create/generate an AppEUI (EUI stands for 'Extended Unique Identifier') and an AppKey. Both these numbers will be needed later in step '6.5 Initializing the flower'.

In this step you should also find your API security key that will be needed in your Node-RED downlink and uplink security settings (see part '5. Node-RED').

For now, that's all you can do. Later you will have to add all your devices (a.k.a. robotic flowers) with their Device EUI (DevEUI) in the application you created. This DevEUI is the unique number defining each LoRa module and can be found by reading it out, but we will come back to this in step '6.5 Initializing the flower'. You will have to create unique names, e.g. flower1, flower2… A tip is to write these names/numbers on the LoRa module or somewhere else on the PCB.

Note that the frequency plan for the radio communication is region bound, e.g. we used Europe 863-870 MHz.

### 4 NODE-RED

Node-Red is a browser based editor needed to receive all the data and creating the daily csv-file on one hand, and on the other hand to generates the dashboard website to monitor all robotic flowers in use. With this programming tool we created a flow of connected JavaScript functions making up the application consisting of a backend and a frontend. The backend receives data from the LoRaWAN backend, processes them and provides them to the frontend application (Node-RED dashboard). The backend also stores the visitation data in a daily CSV-file (an example can be found in supporting information of the paper). For every visit, a new line is added in the file containing the time at the start of the visit, duration of the visit and the name of the device that registered the visit (e.g., 'flower 1'). That

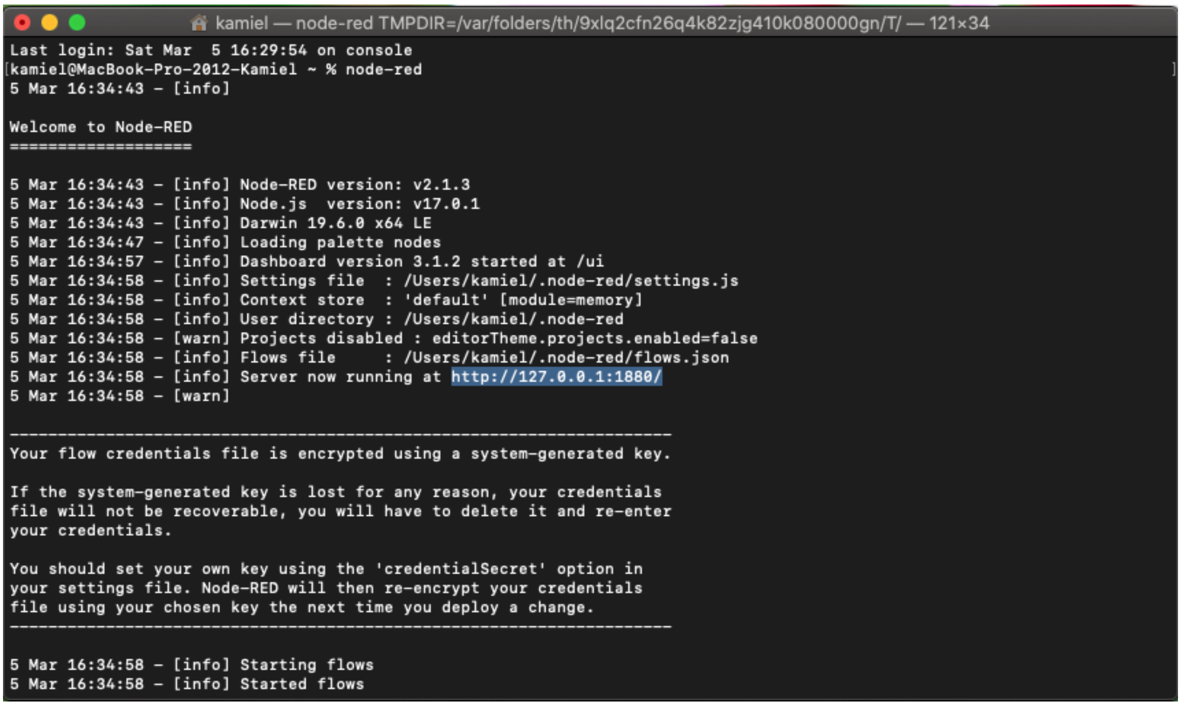

**Figure 11**

is why it is very important to keep Node-RED running continuously when the robotic flowers are in use! If not, data might be lost.

To get started, you need to install Node-RED. For this, there are multiple options (see website of Node-RED: https://nodered.org/ ). You can use a local computer, a device like a Raspberry Pi or have it running in the cloud. We used a computer to run Node-RED locally. Just note that this computer will need to stay running as long as an experiment is running as Node-RED is responsible for all communication from and to the flowers. Different guidelines for installation can be found on the Node-RED website.

After installing on your local machine, you can run Node-RED from the 'Terminal' (macOS) or 'Command Prompt' (Windows) by typing: 'node-red' (Fig. 11). Node-RED will start running and then you can use the displayed server address to go to the web interface. Once you are in the flow editor, you will need get nodes and connections between the nodes in there to build the flows. The program consists of two parts, one for file logging and one for communication with all the robotic flowers separately. These two parts are available in the folder you downloaded from GitHub. Both files are in JSON-format ready to be imported in Node-RED. More information on how to import a flow: [https://nodered.org/docs/user-guide/editor/workspace/import-export](https://nodered.org/docs/user-guide/editor/workspace/import-export).

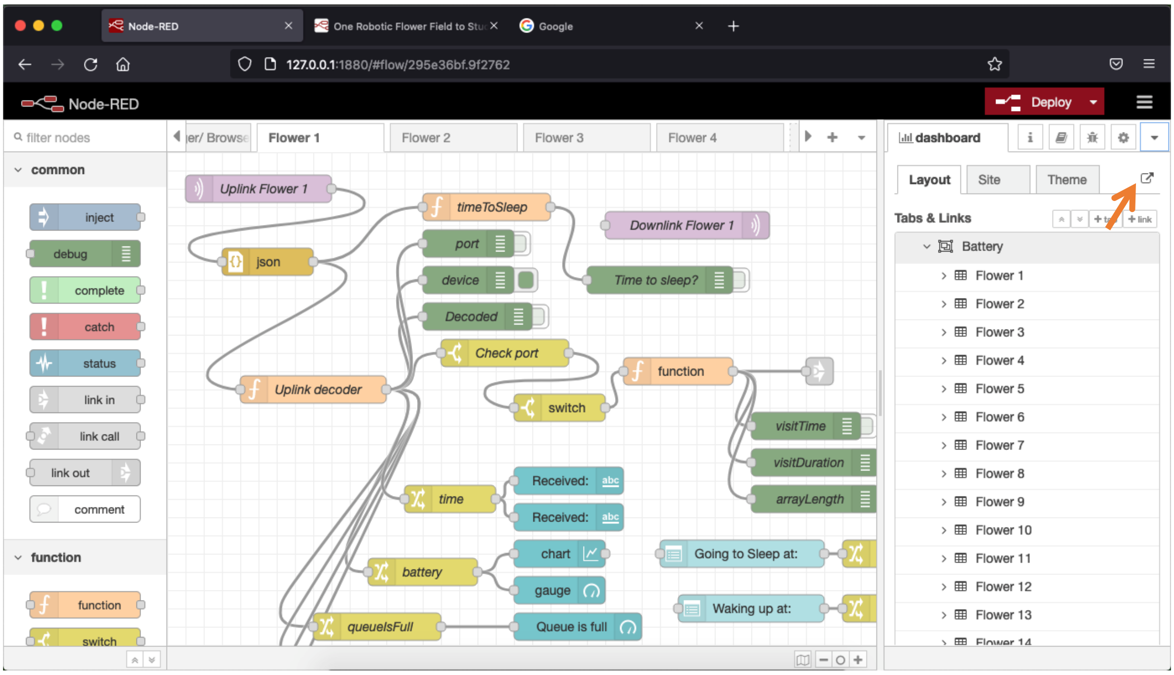

**Figure 12**

If you never used Nod-RED before or don't have the necessary modules installed, you will need to add the following modules to the palette:

- node-red-contrib-fs
- node-red-dashboard
- node-red-node-tail

1. The **'File Logger Flow'** is only needed once per robotic flower field system. Here you need to change the path to where you want a new subfolder 'DataLog' created where your CSV-files will be stored. The nodes where you need to do this are:
    - 'filename generator'
    - 'file-lister'
    - 'init'
    - 'reset'

2. The 'Flower Flow' is needed for every separate robotic flower in the system. So, for example, if you want to use 4 flowers, you will need to import this flow in 4 different tabs (Fig. 12). Also here, you will need to do some important changes for every flower in the Uplink and Downlink nodes:
    - change name and topic so it contains the right device and app name!
    - update server security settings in downlink and uplink node for each flower

So far so good. Now it is time to structure your dashboard website. You can play around with it of course and arrange it as you see fit, but we will show you how we did it (Fig. 13). First go to the dashboard page, opening it on the left side of your screen as can be seen in Fig.12. There, we created the different tabs for the dashboard: Battery, Alarms, Sleep time, Refill gap and Files. Next, in each tab except 'Files', we made a group for each flower.

We constructed the dashboard in such a way that, for example, the 'battery' tab shows an overview of all the robotic flowers that are in use with the time of the last update (last received message), the battery voltage, and a graph of the voltage decline in function of time. We assigned the the necessary nodes per flower to the right tab and group, an example for flower 1 given below:

- Received: '[Alarms] Flower1'
- Received: '[Battery] Flower1'
- Chart:'[Battery] Flower1'
- Gauge: '[Battery] Flower1'
- Queue is full: '[Alarms] Flower1'
- Send during visit: '[Alarms] Flower1'
- Battery warning: '[Alarms] Flower1'
- Possible obstruction: '[Alarms] Flower1'
- Refill gap: '[Refill gap] Flower1'
- Going to sleep at: '[Sleep time] Flower1'
- Waking up at: '[Sleep time] Flower1'

One can also see the benefit of having a Tab for each flower (e.g., could be useful for use on smartphone), but here we like the quick overview of all the flowers in one tab.

To see the dashboard website you just created, click on the little link symbol in the top (Fig. 12, arrow). In 'Theme' you can later customize colours, titles etc. of your dashboard website.

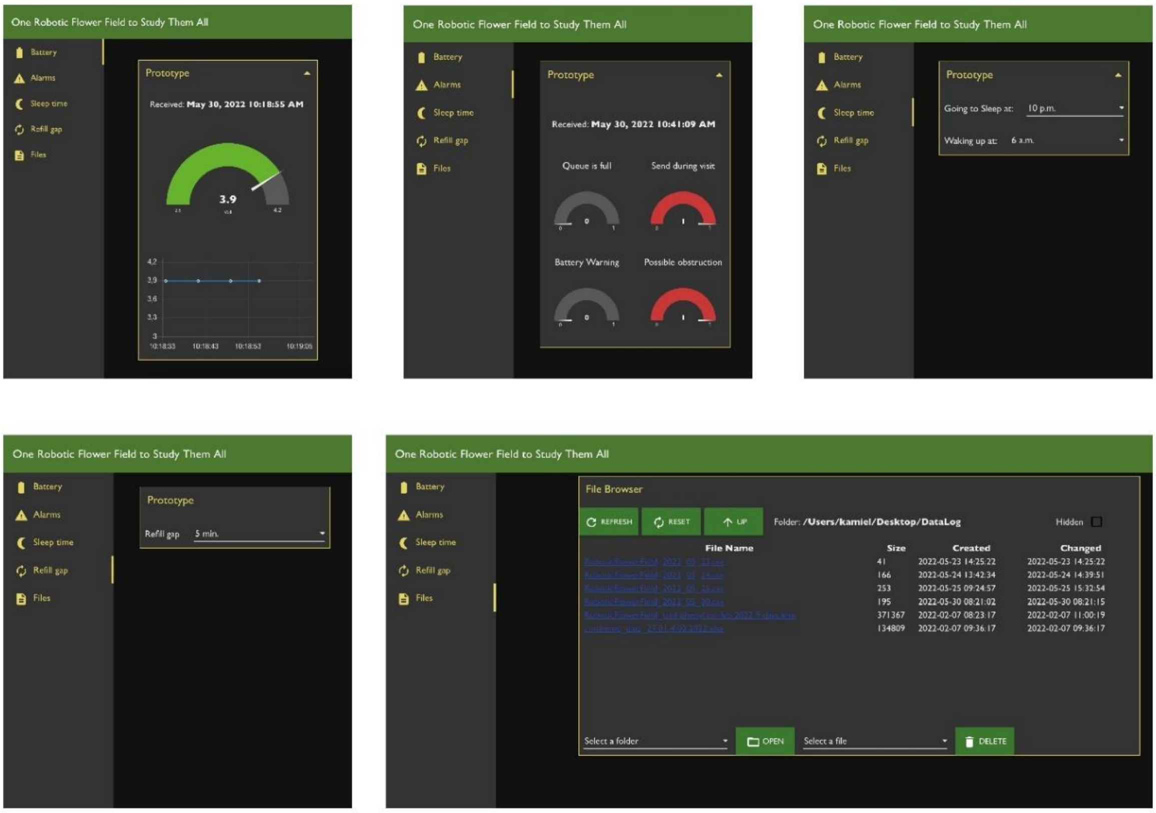

**Figure 13**

### 5 UPLOADING FIRMWARE TO ARDUINO

#### 5.1 SOFTWARE INSTALLING

To be able to see, edit and upload the firmware, a code editing software is needed. We used Visual Studio Code (VSC) as it is a free program that is built on open source and offers great flexibility. You are of course welcome to use whatever other editor you might have experience with.

To get started, you will need to download Visual Studio Code from https://code.visualstudio.com/

Next, you can install PlatformIO from the extension list when you have opened the freshly installed VSC (Fig. 14). PlatformIO is an integrated development environment (IDE) that we have experienced as very well suited for this kind of project. You can also already install C/C++ extension pack, or do it later when VSC prompts you to do so.

#### 5.2 DOWNLOAD FIRMWARE

Remember the folder you saved before from GitHub? Now, we will need to do some stuff there before continuing. In the folder 'Firmware', you can find the subfolder 'src' in which 2 files can be found each containing a slightly different version of the firmware controlling the robotic flower. Before you can proceed to the next steps, you will have to remove one of both out of the firmware folder as you can only use one of both at once.

Choose the firmware file best suited for your needs:

1) 'main.cpp' has automatic refilling and a refill with refill gap (that can be chosen from the Node-RED dashboard website) after each visit. E.g., on top of the automatic refilling every 10 minutes (or other time specified when uploading the firmware) to prevent drying out, the flower will also refill after a bee visits (leaving a gap time that can be chosen remotely).

2) 'main_autoRefillOnly.cpp' only has the automatic refilling (that can be chosen from the Node-RED dashboard website remotely)

Now you can open the firmware folder using VSC. Once you opened the files in the editor, you can use the explorer on the right of the screen (Fig. 14) to see all the files.

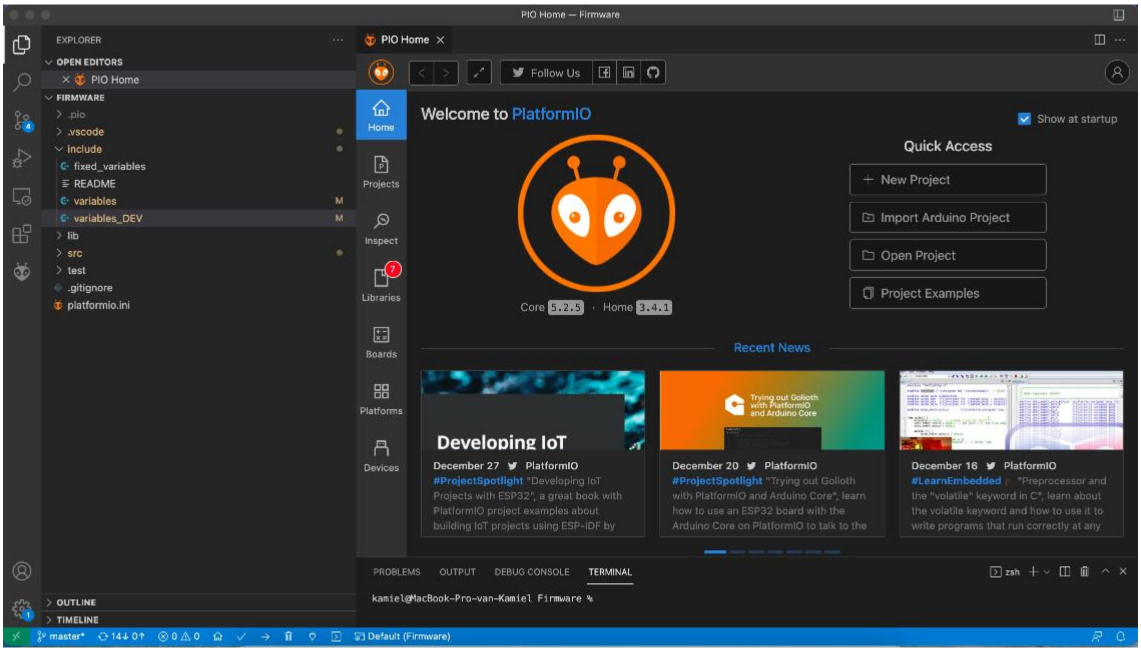

**Figure 14**

#### 5.3 Modify firmware to fit needs

If you want to change parameters so they fit your exact research questions, you can open the file ‘variables’ in the folder ‘include’. Here you can find all the parameters with some explanation and easily change them before uploading to the Arduino (Fig. 15).

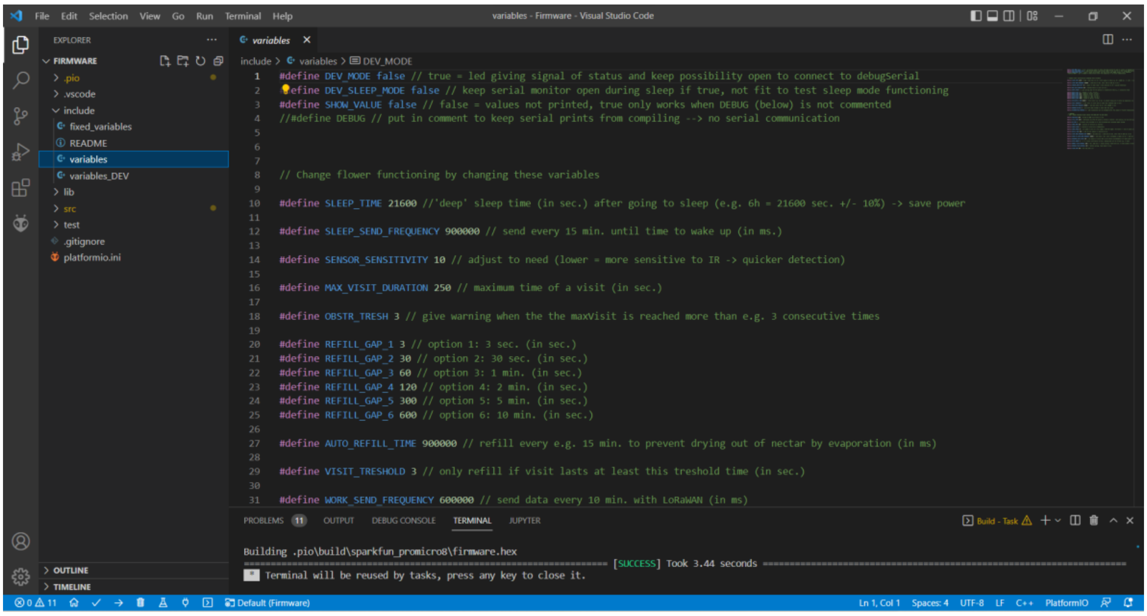

**Figure 15**

##### 5.3.1 *List of adjustable parameters*

- **SLEEP_TIME** 21600 *// 'deep' sleep time (in sec.) after going to sleep (e.g. 6h = 21600 sec. +/- 10%) -> save power*
- **SLEEP_SEND_FREQUENCY** 900000 *// send every 15 min. until time to wake up (in ms.)*
- **SENSOR_SENSITIVITY** 10 *// adjust to need (lower = more sensitive to IR -> quicker detection)*
- **MAX_VISIT_DURATION** 250 *// maximum time of a visit (in sec.)*
- **OBSTR_TRESH** 3 *// give warning when the the maxVisit is reached more than e.g. 3 consecutive times*

- **REFILL_GAP_1** 3 *// option 1: 3 sec. (in sec.)*
- **REFILL_GAP_2** 30 *// option 2: 30 sec. (in sec.)*
- **REFILL_GAP_3** 60 *// option 3: 1 min. (in sec.)*
- **REFILL_GAP_4** 120 *// option 4: 2 min. (in sec.)*
- **REFILL_GAP_5** 300 *// option 5: 5 min. (in sec.)*
- **REFILL_GAP_6** 600 *// option 6: 10 min. (in sec.)*

- **AUTO_REFILL_TIME** 900000 *// refill every e.g. 15 min. to prevent drying out of nectar by evaporation (in ms)*
- **VISIT_TRESHOLD** 3 *// only refill if visit lasts at least this treshold time (in sec.)*
- **WORK_SEND_FREQUENCY** 600000 *// send data every 10 min. with LoRaWAN (in ms)*
- **WARNING_VOLTAGE** 3 *// give alarm when voltage is lower than this value*
- **RECONNECTION_TRESHOLD** 2 *// LoRa reconnection will be attempted after this amount of failed transmissions*

#### 5.4 Upload to Arduino

Now you are ready to upload the firmware to your Arduino Pro Micro. To do that, you will have to connect the Arduino with a USB-cable to your computer. Next, navigate to the ‘main.cpp’ file (depending on which one you chose earlier on) in the ‘src’ folder.

Once you are in this file, the first thing you need to know is which variables to include in line 11 (Fig.16). We created a second set of variables called ‘variables_DEV’ that give access to a ‘development mode’ in the Robotic Flower (see section 7.1). This is made so you can more easily check the functioning of the flower and track any problems if needed, e.g. by faster sending times, keeping the serial monitor open and a LED signalling the state of the flower. To include ‘variables’ or ‘variables_DEV’, you will have to change this in the file by writing the name of the chosen one.

To build the firmware, press the little ‘√’ in the bottom right, next to upload the built firmware to the Arduino press the ‘→’-button (Fig.16). When uploading is successful, the terminal will tell you so and then you can open the serial monitor with the ‘ ’-button.

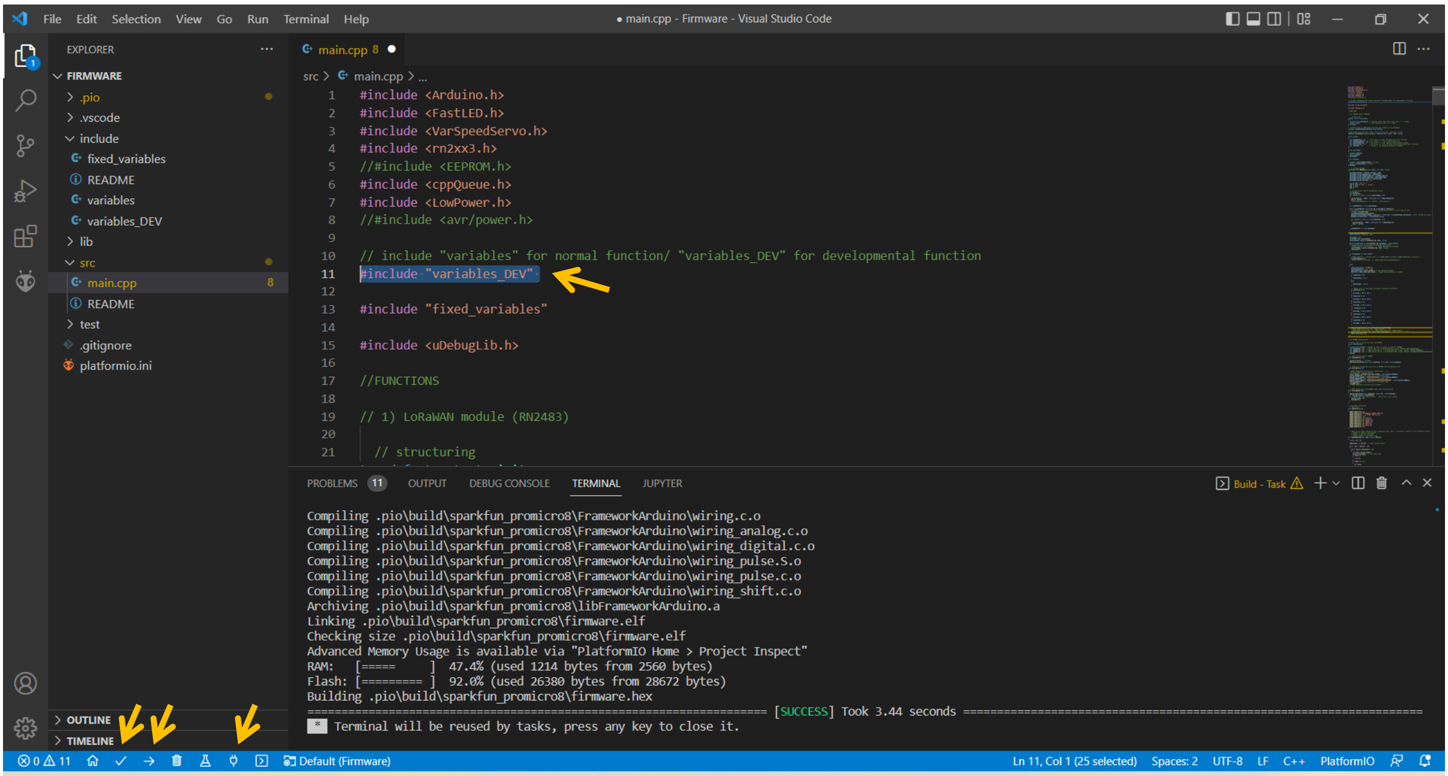

**Figure 16**

#### 5.5 INITIALIZING THE FLOWER

The first time a new robotic flower is connected, the parameters that are needed to make connection with an IoT network must be stored in the permanent memory of the microcontroller. To do this, upload the firmware in development mode, i.e. with including the ‘variables_DEV’, and open the serial monitor with the ‘ ’-button in the bottom left (see Fig. 16). After some seconds, a menu will show up (Fig. 17). When you type 1 in the command line and press enter, the serial monitor will show the current parameters (initialized at all 0’s when not entered before). Here you will be able to find the DevEUI you need to complete part ‘4. Setup IoT’, where devices should be added to the application. By typing 2 in the menu, the DevEUI can be altered manually, but this should not be necessary. Typing 3 and 4 respectively in the menu lets you configure the AppEUI and AppKey (see part ‘4. Setup IoT’).

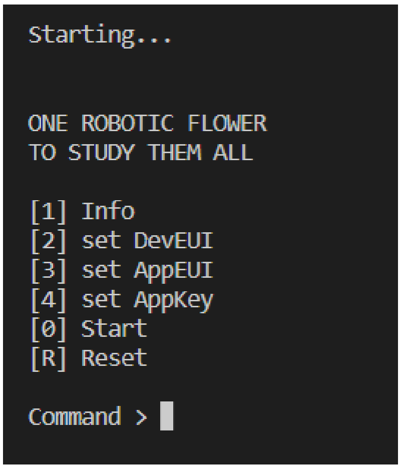

**Figure 17**

When all these parameters have been set you can type 0 in the command line to start the functioning of the flower for real. Moreover, when switching on battery power, you will also see the LED light signalling the functioning and the movement of the servo arm (we recommend not attaching the servo arm yet, as the position of the servo needs to be initialized first, see ‘7.1 Preparations’).

##### *5.5.1 SERIAL MONITOR*

In development mode, you can follow what is happening on the serial monitor in your computer. With prints coming from the Arduino through the USB-cable, you can see where in the firmware the Arduino is operating.

If you want or don’t want to see the reading of the IR-sensor in the serial monitor during development mode, you can specify this in the top of the ‘variables_DEV’ file in the folder ‘Include’ in VSC. Putting True or False after ‘SHOW_VALUE’ will do the trick. Seeing the values can help you make sure the detection system works as it should be: put something like a pencil inside the feeding hole and you should see a significant drop in the IR-value. In ours, the baseline (no visit) value is approx. 950.

To more easily test the flower with sleep mode, there also is a variable ‘DEV_SLEEP_MODE’ that can be on True or False in the ‘variables_DEV’ file. This is because the regular sleep mode puts the Arduino in a power saving state which automatically breaks of the serial connection, meaning that the serial monitor will stop working.

##### *5.5.2 LED SIGNALS*

The LED only works when the battery is switched on (just like the servo motor the LED works on 5V, unlike the voltage coming through the Arduino which is 3.3V only).

Led signals during starting up: begins with red, it goes out when looking for serial monitor, red again when attempting to make connection and finally blinking green twice quickly when connection is established. Then the flower starts working.

- Blinking green every second = working mode, battery normal
- Blinking red every half second = working mode, battery warning (can become very dim when battery almost empty)
- Continuously red = detecting a visit
- Flashing bright blue = end of visit and visit saved
- Continuously yellow = sending data over LoRaWAN
- Continuously deep blue = in sleep mode

##### 5.5.3 *OPERATIONAL MODE*

When putting flower in operational mode by including ‘variables’ in stead of ‘variables_DEV’, the flower will not longer show LED signals after connection is made and the serial monitor will be closed together with the USB communication on the Arduino to save battery power.

Once the program is on the Arduino, it will start running from the moment when the battery is powered on, so opening the serial monitor with the USB-cable etc. is not longer necessary. If no serial connection is detected, the flower automatically proceeds to working.

!! Note that, to upload new variables or put the flower back in development mode, you will need to take away all the power sources to the Arduino (battery and USB) before connecting the USB again and upload the firmware again before the USB communication is blocked when the flower is starting up.

### 6 STEPS TO PREPARE A TRIAL

#### 6.1 PREPARATIONS

Now that we have our flowers ready to collect data, we will need to set them up in the correct way to make them work as long as we want.

1. First, we will need to get the batteries charged. A fully charged battery can power a robotic flower for approximately two weeks (see part ‘8. Battery life estimation’). You can charge the batteries using the battery module with the micro USB port, but it can also be convenient to have a separate charger. In this way you can have spare batteries charging while the flowers are working.

2. Then, it can be useful later to have all the flowers labelled on the outside with a sticker (or in another way) with the same device name you put in the application before (part ‘4. Setup IoT’). For example, label them with numbers, so that you easily can see which flower is which during the trial.

3. Make sure all parts are clean and dry. Especially if they have been used before, make sure all leftover artificial nectar is cleaned. Watch out, as 3D printed parts are not suited for the dishwasher! Best is to just use semi-hot water and soap to clean these parts. See in ‘9. Troubleshooting’ the part about cleaning PCB.

4. Probably you will want to offer some kind of reward to the little visitors in the robotic flower. You can make any solution you need for your experiment and fill the nectar reservoir. It is best to fill it with at least 40 ml, but you can go up to 80 ml (that is, if you use the same container we did; see also part ‘3.3.5 Nectar reservoir’). But don’t put it inside the flower yet, as the flowers will need to be moved. To prevent any spilling, we place it in the nectar reservoir case (see part ‘3.3.4 Reservoir case’) only as a last step of the flower setup.

#### 6.2 ATTACHING SERVO ARM

Something we didn’t do yet is attaching the servo arm. See Fig. 8 for an example of an attached servo arm. The first step is to attach the steel wire to the nectar cup (see part ‘3.3.3 Nectar cup’) at one side and the spindle on the other side. While doing this, keep in mind the distance between the nectar hole and the servo motor to choose the length of your steel wire (better to take it a little bit too long than too short). When this is done, we can now start to attach it to the servo motor in the right way as to ensure good alignment with the nectar hole.

1. The start is to get the right distance and angle in the steel wire. Attach the servo arm to the servomotor by pushing the spindle on the gear, as if it would be in closed position. Now you can bend the steel wire until the nectar cup fits nicely the notch in the central flower disk. You can check the nectar cup alignment with the feeding hole by looking from above. When this step is done, take the servo arm of again.

2. Next, make sure the servo is in closed position. To do this, the flower needs to be working on battery power. The easiest ways to proceed from here is to put the flower in development mode, or have the refilling rate at e.g. 30 seconds via Node-RED. Now wait until you hear the flower refilling at least once, so that you know that the servo is in closed position.

3. Now you can press the spindle on the gear of the servomotor, it doesn't matter in which position. Then forcibly turn the gear a (little) bit up and take the arm out again. Then attach the servo arm in the right position (touching the hole). This step ensures a tight fit, as the servo arm will be kind of 'pressed' against the nectar hole.

4. Now check again if the position is still correct after a couple of servo movements. If not, carefully adjust the steel wire to correct the nectar cup position.

#### 6.3 Experimental Setup

1. The first step: choose the right firmware variables for your experiment. See part '6.3.1 List of adjustable parameters' to see all the options. Be critical when setting these, also paying attention if the value e.g. is given in seconds or milliseconds. When ready, upload to the Arduino. If nothing has to change since the last used firmware, you can skip this part.

2. Put flowers together:

   - Make sure all cables are connected and connectors are plugged in.
   - Put the PCB inside the stem, making sure the holes fit the protrusions on the bottom.
   - Next, put the battery on top of the PCB in horizontal position, making sure it stays low
   - Insert the reservoir case, making sure it fits the protrusions in the stem and the cables going to the servo are at one side in the cable hole and the cables going to the IR-lights are in the other side. The X-marking on the reservoir case should be at the side where the servo arm will go down in the nectar reservoir! Put the reservoir case low enough so that is rests on the support beams in the stem.
   - Now you can put the background plane with central flower disk on top and close the robotic flower with the twist-and-lock system.
   - The nectar reservoir with artificial nectar can be added later, when flowers are in position.

3. Start Node-RED and open the flow website in your browser. Now you have the option to double click the flow tab for each flower to enable or disable it, e.g. if you're only using a subset. Disabled flowers will disappear from your dashboard website, so they don't distract you when monitoring the flowers in use.

4. Important to keep in mind when using Node-RED on a local computer is that you let it run during experiment. This means the computer itself needs to stay on (not in sleep mode), the terminal/ command prompt has to stay open and internet connection should not be disrupted.

5. Now open the dashboard website and set the parameters as you want them:

    - Sleep times, ‘Going to sleep at’ and ‘Waking up at’, are not mandatory to set, but will increase the battery life-time. Take in mind the flower will always be sleeping for at least the period of ‘SLEEP_TIME’ defined in the variables. We often set use sleeping from 22 pm to 6 am, with 6h of deep sleep.

    - Setting the Refill gap is mandatory. You can choose between the different options you set before (names can be adapted in the Node-RED flow to match the ones you set in ‘Variables’).

6. Set your robotic flowers in the formation and where you want, for example setting them up in patches (blue vs yellow flowers containing nectar of different composition) in a greenhouse (Fig. 18). Tip: make a scheme of the setup with flower numbers and their different treatments.

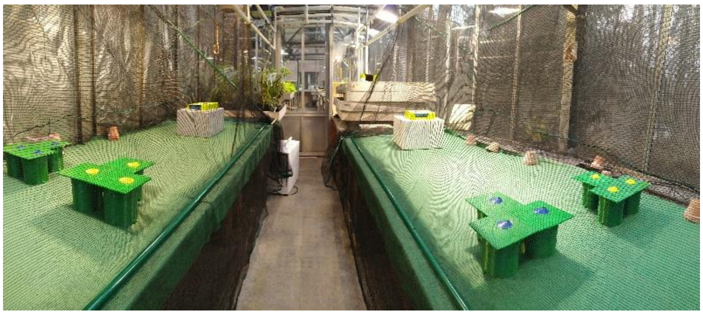

**Figure 18**

7. Put the nectar reservoir in the reservoir cage inside the flower and close again. Check if the X-mark on the reservoir cage matches the side of the feeding hole.

8. Switch the flowers on with the switch.

9. Go to dashboard to see if all flowers are connected and working. An easy way to do this is to click battery status and check if all flowers have been sending.

10. Release the study animals and let the data come in!

11. Bonus tip: it can be favourable to start with a learning phase, where the study animals are allowed to discover the robotic flowers in a smaller arena (Fig. 19).

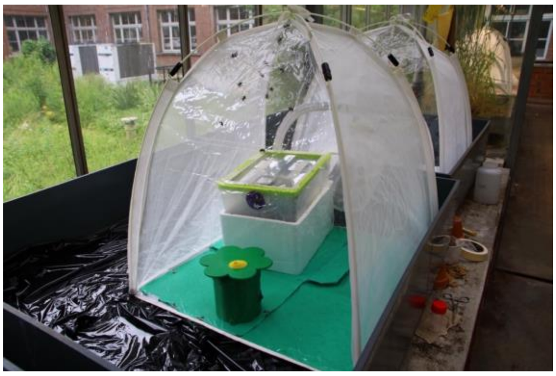

**Figure 19**

#### 6.4 DURING TRIAL

A daily CSV file will be created in a folder called ‘DataLog’ on the location you have specified in Node-RED ‘file logger flow’ (see part ‘5. Node-RED). Data is added to the bottom of the daily file in real-time. If you change the name or the location of the file of the day while data is still coming in, a new will be started in the ‘DataLog’ folder. Each flower will send its gathered data every 10 minutes. In between sending, these data are saved in RAM, meaning that they are lost when suddenly power is down!

##### *6.4.1 DASHBOARD MONITORING*

During the trial you can keep an eye on the functioning of the flowers by looking at the dashboard website (Fig. 13).

Using the ‘Battery’ tab, you can check the last time data has been sent and the battery level of the flowers. To have an idea how long the flowers can keep functioning with the current voltage level, you can compare with the graph in ‘Fig. 22. Battery voltage over time’. If needed, batteries can be changed during the trial. Be aware though that it might cause some loss of data stored in RAM when the flower loses power. A solution is to do this battery change after going to sleep time. Or you can take the flower out of the arena carefully and connect the battery module to power with micro-USB while changing the battery. This way power continuity is ensured.

In the tab ‘Alarms’ you can check if there are any other problems popping up:

- Queue is full, meaning the flowers RAM is full and no more visits can be stored. This means the flower is getting visits at a higher rate then it can send. Every 10 minutes, data of 16 visits can be send, with a buffer of 100 visits.
- Sending during visit.
- Battery warning, meaning the battery is lower than the specified threshold.
- A possible obstruction of the IR-sensor.

Using a server or running Node-RED in the cloud you can access the dashboard everywhere easily. But what if you are using a local machine for running Node-RED, e.g. on a computer in the lab as we did, but you are at home and want to check out the dashboard? That is possible by installing Node-RED for example on your personal laptop, but here you only put the flower flow (not ‘file logger flow’) and make sure you delete the uplink nodes, so that the flowers only get instructions from the lab computer.

### 7 BATTERY LIFE ESTIMATION

#### 7.1 THEORY

A rechargeable 18650 Li-Ion 3.7 V button-top battery with 3400 mAh (VABO NV, Heusden-Zolder) was used. The number 18650 is based on the size of the battery, which is 18 mm diameter and 65 mm in length.

The batteries have a built-in protection circuit that protects the batteries from overcharging and over-discharging; both could indeed damage the cell and possibly even lead to a safety hazard (fire and/or explosion). Li-Ion cells should be handled with care.

The 3.7 V gives the nominal or average voltage. A fully charged battery will be maximum 4.2 V and in use the voltage will drop until the minimum is reached (Fig. 1; Chen & Rincon-Mora, 2006). Charge time is approximately 5.5 hours according to the manufacturer data sheet.

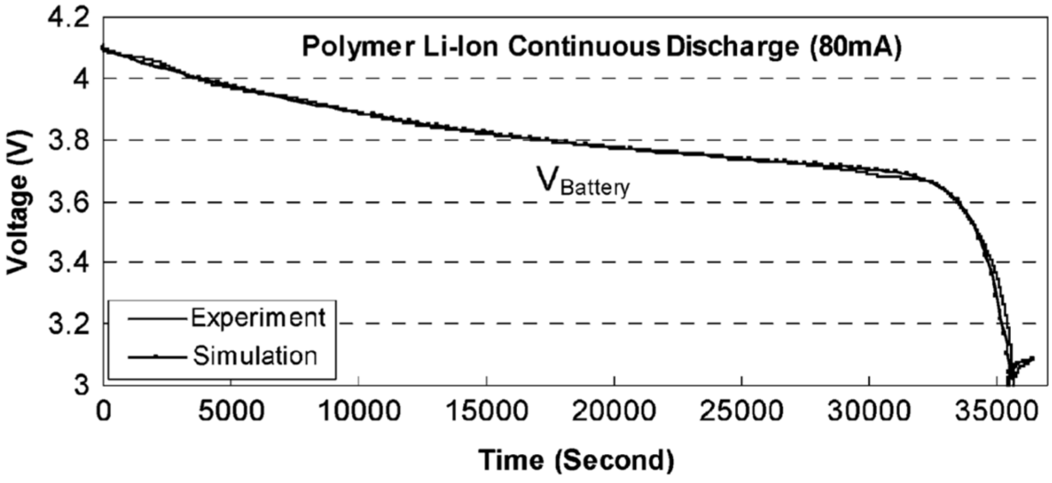

**Figure 20:** ***A typical voltage response curve, showing the maximum voltage around 4.2, the minimum at 3 and the nominal voltage around 3.7 for lithium-ion batteries (Chen & Rincon-Mora, 2006).***

The 3.7 V Li-Ion batteries that were used here have 3400 mAh (milliampere hour). This value represents the total amount of energy that can be stored in the battery, which combined with the mean energy consumption can give an indication of how long the battery will be able to power the robotic flower.

First, we need to take the conversion of 3.7 V to 5 V in the battery module into account. As the wattage ($W = V \times I$, with V in volts and I in ampere) before and after conversion stays the same, the formula $I_2 = {}^{V_1}/_{V_2} \times I_1$ can be used to determine the new value for mAh. In this case, that gives 2516 mAh, but there is a loss of energy during the conversion that must be considered. By taking a (wide) margin of 10% loss, the value used for further calculations of battery life is 2264 mAh.

Next, the energy consumption of the robotic flower needs to be known. For this, we performed a 24-hour measurement, containing the work state and the sleep state, to find the mean value for energy consumption of the robotic flower. This was done using JoulescopeTM, a precision energy analyser that can be connected to a computer and, among others, show changes over time like an oscilloscope (Fig. 2). JoulescopeTM was also used during software development to reduce energy consumption. Using parameters 10 pm and 6 am for going to sleep and waking up time respectively with 7 hours of deep sleep and refilling rate as well as sending rate at 10 minutes, the mean value for basal energy consumption (without visitors) over 24 hours was 4.1 mA. This value combined with the battery capacity yields a predicted battery lifetime of 19.56 days according to the Oregon Embedded online battery life calculator[1] (which reduces the lifetime with 15% automatically to account for self-discharge of the battery).

---

[1] https://oregonembedded.com/batterycalc.htm

This value is an estimation, as several factors might influence the battery life, one of them being the ambient temperature in which the robotic flower is used. Li-Ion batteries usually have an optimal operating temperature between 15°C and 35°C, higher or lower temperatures may affect the battery performance negatively (Ma et al., 2018). For example, when using the robotic flowers in a glasshouse, it is good to know that high temperatures could lead to a reduced lifetime of the batteries.

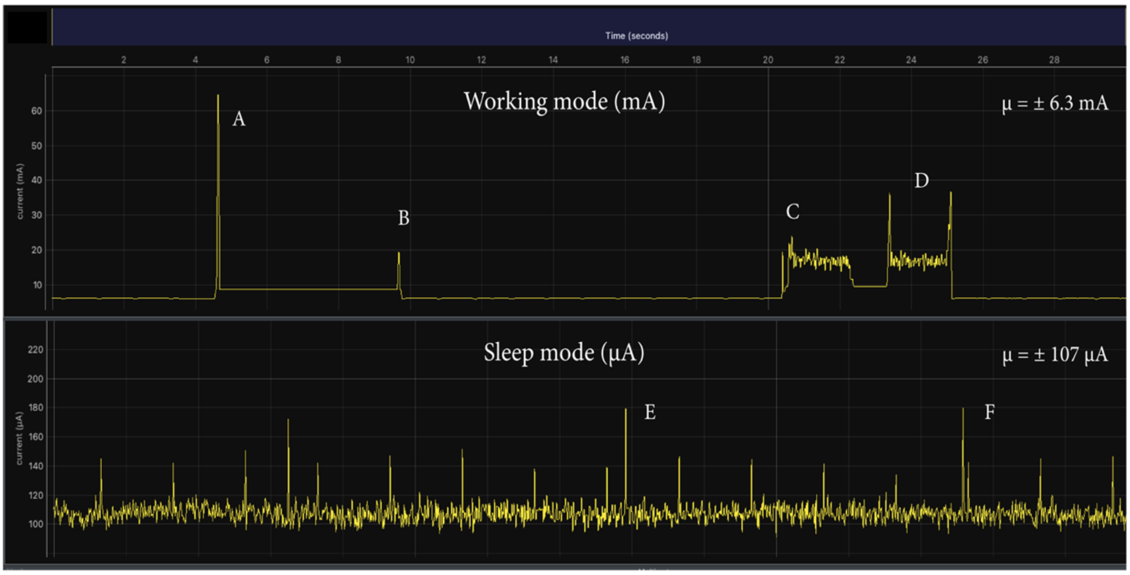

**Figure 21:** ***Energy consumption of the robotic flower during working mode and sleep mode with A) the spike of the LoRaWAN module sending data, B) the LoRaWAN module receiving a message after sending, C) the servomotor moving to the open position, D) the servomotor moving to the close position, E) and F) showing two energy consumption spikes during sleep mode. Made with JoulescopeTM.***

The mean energy consumption during the work state was around 6.3 mA without floral visitation (basal energy consumption). This value went up to 6.8 mA when a visit of variable length (but longer than the visit threshold of three seconds) was simulated every minute, which is logical as the servomotor will have to work more frequently and the LoRaWAN module will have to send larger data packages each time.

#### 7.2 IN PRACTICE

To test the actual basal (i.e., without pollinators visiting) battery lifetime, we set-up a flower with a fully charged battery (± 4.1V) in an office with a relatively constant temperature around 20 °C. The sleep time was set between 10pm and 6am. For each day, the voltage level was measured 12 times until the flower stopped (± 2.5V). These data are represented in Fig.3, showing the voltage over time. The result was a lifetime of 15 days, but to avoid loss of data because of empty batteries we recommend a 14-day limit. The graph can be used to predict the remaining lifetime of the battery for a given voltage level.

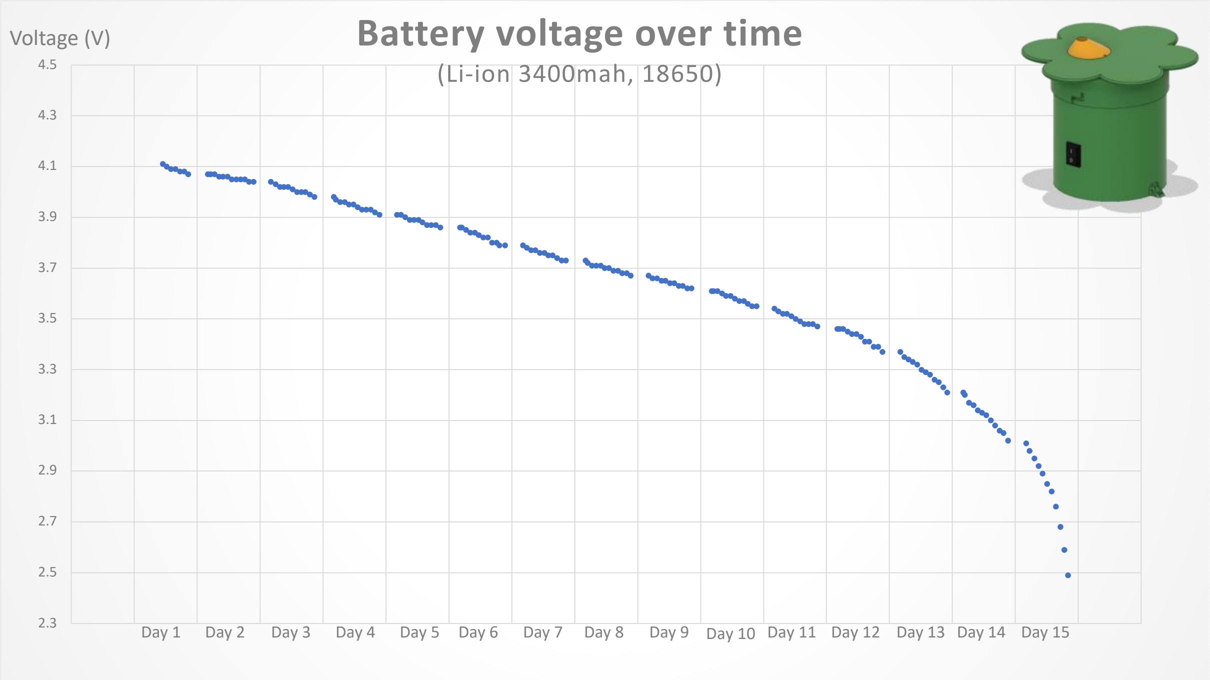

**Figure 22:** ***Evolution of the robotic flower battery voltage over time.***

### 8 TROUBLESHOOTING

To check the well-functioning of the robotic flowers, we recommend you to play around with them in the development mode. Here you can follow everything that is happening in the serial monitor. Before use, check if the servo motor and IR detection system is working. If you find a problem, try to deduce what might be the cause by ruling things out step by step. Here we list some things that might happen to your flowers with some hints that might help you solve the problem. If problems persist, feel free to contact us for help!

1. You don't see the flowers in Node-RED dashboard

   ➔ Make sure the flowers you want to use are enabled in the Node-RED flow overview.

2. You get a spill on the PCB and want to clean it

   ➔ Spills on the PCB can lead to small problems, e.g. misreading of battery level, but also to serious malfunctioning. You can clean this, but always use demineralized water to avoid crystals after drying. We had this problem and fixed it with emerging the PCB in a supersonic bath of demineralized water at 50°C for 5 minutes (without antenna), gently brushing it with a toothbrush. It is very important to let it dry thoroughly before using it again, we used a hot air oven at 50°C for this purpose.

3. A flower did not wake up from sleep mode

   ➔ Make sure you put the correct sleeping times in Node-RED. By switching the flower off and on again it should reset and start working again. If not, try changing the battery. If the problem exists in all the flowers, check the firmware for any mistakes, e.g. in the variable 'SLEEP_TIME'.

4. The flower is working, but is seems to be stuck in trying to make connection

   - ➔ Check if Node-RED is running and if all parameters are set correctly in the IoT service you use (e.g. The things Network) and correspond to the parameters set in the flower. Does the flower show the LoRa module version and DevEUI when typing 1 in the start menu in Development mode (see part '5.5 Initializing the flower')? If not, the problem might be due to bad connection of the LoRa module on the PCB. Check if soldering was done correctly and/or if any connections have been made where there should not be. Cleaning the PCB might also help (see Troubleshooting 3).

5. The flower has suddenly stopped sending data

   - ➔ Check if the battery is still charged. If that is not the problem, it might be the battery module that is failing. Switching the flower off and on again can reset the flower. If this does not help, check for any hardware problems: is there a cable not connected anymore? Is there a big spill on electronics?

6. The flower does not detect any visits

   - ➔ Put the flower in development mode and check the IR values the flower is receiving. You should see a drop in this value when you manually block the sensor. If nor, it might be the sensor or emitter is broken or not connected correctly.

7. The flower does not start working and the battery module does not show any LED light when switched on

   - ➔ First check if the battery is charged and inserted correctly (+ and -). If that is not the problem, you can try to reset the battery module by connecting it shortly with the micro-USB port to a power source. If this is still not working, try with another battery module.

### 9 REFERENCES

Balfour, N. J., Garbuzov, M., & Ratnieks, F. L. W. (2013). Longer tongues and swifter handling: why do more bumble bees (Bombus spp.) than honey bees (Apis mellifera) forage on lavender (Lavandula spp.)? *ECOLOGICAL ENTOMOLOGY*, *38*(4), 323–329. https://doi.org/10.1111/een.12019

Branquart, E., & Hemptinne, J. L. (2000). Selectivity in the exploitation of floral resources by hoverflies (Diptera : Syrphinae). *ECOGRAPHY*, *23*(6), 732–742. https://doi.org/10.1034/j.1600-0587.2000.230610.x

Chen, M., & Rincon-Mora, G. A. (2006). Accurate electrical battery model capable of predicting, runtime and I-V performance. *IEEE TRANSACTIONS ON ENERGY CONVERSION*, *21*(2), 504–511. https://doi.org/10.1109/TEC.2006.874229

Corbet, S. A. (2000). Butterfly nectaring flowers: butterfly morphology and flower form. *ENTOMOLOGIA EXPERIMENTALIS ET APPLICATA*, *96*(3), 289–298. https://doi.org/10.1046/j.1570-7458.2000.00708.x

Kuusela, E., & Lämsä, J. (2016). A low-cost, computer-controlled robotic flower system for behavioral experiments. *ECOLOGY AND EVOLUTION*, *6*(8), 2594–2600. https://doi.org/10.1002/ece3.2062

Ma, S., Jiang, M., Tao, P., Song, C., Wu, J., Wang, J., … Shang, W. (2018). Temperature effect and thermal impact in lithium-ion batteries: A review. *Progress in Natural Science: Materials International*, *28*(6), 653–666. https://doi.org/https://doi.org/10.1016/j.pnsc.2018.11.002

Pozo, M. I., van Kemenade, G., van Oystaeyen, A., Aledon-Catala, T., Benavente, A., den Ende, W., … Jacquemyn, H. (2020). The impact of yeast presence in nectar on bumble bee behavior and fitness. *ECOLOGICAL MONOGRAPHS*, *90*(1). https://doi.org/10.1002/ecm.1393

### 10 Appendix

#### 10.1 Parts List

| Part | Specifications | Article number | #/flower | Company | URL |
|---|---|---|---|---|---|
| **Microcontroller Pro Micro** | ATMEGA32U4 3.3V/8MHZ | DEV-12587 | 1 | **DigiKey** | https://www.digikey.be/product-detail/en/sparkfun-electronics/DEV-12587/1568-1061-ND/5140826 |
| **Micro Servo** | SERVOMOTOR RC 4.8V | SER0006 | 1 | **DigiKey** | https://www.digikey.be/product-detail/en/dfrobot/SER0006/1738-1385-ND/7597224 |
| **LED (per 5)** | ADDRESS LED DISCR SERIAL RGB 5PC | COM-12986 | 0.2 | **DigiKey** | https://www.digikey.be/product-detail/en/sparkfun-electronics/COM-12986/1568-1213-ND/5673799 |
| **ON/OFF Switch** | SWITCH ROCKER SPST 10A 125V | RA1113112R | 1 | **DigiKey** | https://www.digikey.be/product-detail/en/e-switch/RA1113112R/EG5619-ND/3778055 |
| **IR emitter** | EMITTER IR 940NM 50MA RADIAL | IR908-7C-F | 1 | **DigiKey** | https://www.digikey.be/product-detail/en/everlight-electronics-co-ltd/IR908-7C-F/1080-1089-ND/2675580 |
| **IR receiver** | SENSOR PHOTO 940NM SIDE VIEW RAD | PT908-7C-F | 1 | **DigiKey** | https://www.digikey.be/product-detail/en/everlight-electronics-co-ltd/PT908-7C-F/1080-1164-ND/2675655 |
| **Power Switch** | IC PWR SWITCH N-CHANNEL 1:1 8DIP | TPS2024P | 1 | **DigiKey** | https://www.digikey.be/product-detail/en/texas-instruments/TPS2024P/296-12234-5-ND/413526 |
| **Level Converter** | IC BUF NON-INVERT 5.5V 14DIP | SN74HCT125N | 1 | **DigiKey** | https://www.digikey.be/product-detail/en/texas-instruments/SN74HCT125N/296-8386-5-ND/376860 |
| **Diode** | DIODE SCHOTTKY 30V 1A DO41 | 1N5818 | 1 | **DigiKey** | https://www.digikey.be/product-detail/en/stmicroelectronics/1N5818/497-4548-1-ND/770973 |
| **Res 200** | RES 200 OHM 1/2W 1% AXIAL | SFR16S0002000FR500 | 2 | **DigiKey** | https://www.digikey.be/product-detail/en/vishay-beyschlag-draloric-bc-components/SFR16S0002000FR500/PPC200XCT-ND/594751 |
| **Res 2k** | RES 2K OHM 1/2W 1% AXIAL | SFR16S0002001FR500 | 2 | **DigiKey** | https://www.digikey.be/product-detail/en/vishay-beyschlag-draloric-bc-components/SFR16S0002001FR500/PPC2.00KXCT-ND/594777 |
| **Res 10 k** | RES 10K OHM 1/2W 1% AXIAL | SFR16S0001002FR500 | 3 | **DigiKey** | https://www.digikey.be/product-detail/en/vishay-beyschlag-draloric-bc-components/SFR16S0001002FR500/PPC10.0KXCT-ND/594695 |
| **Res 100k** | RES 100K OHM 0.4W 1% AXIAL | MBA02040C1003FRP00 | 2 | **DigiKey** | https://www.digikey.be/product-detail/en/vishay-beyschlag-draloric-bc-components/MBA02040C1003FRP00/BC100KXCT-ND/336963 |
| **Capacitor 100nF** | CAP CER 0.1UF 50V X7R RADIAL | C322C104K5R5TA7301 | 8 | **DigiKey** | https://www.digikey.be/product-detail/en/kemet/C322C104K5R5TA7301/399-9877-1-ND/3726183 |
| **Capacitor 33uF** | CAP ALUM POLY 33UF 20% 20V T/H | RNS1D330MDS1 | 2 | **DigiKey** | https://www.digikey.be/product-detail/en/nichicon/RNS1D330MDS1/493-14237-ND/4991537 |
| **Antenna** | RF ANT 868MHZ WHIP STR SMA MALE | ANT-868-CW-RH-SMA | 1 | **DigiKey** | https://www.digikey.be/product-detail/en/linx-technologies-inc/ANT-868-CW-RH-SMA/ANT-868-CW-RH-SMA-ND/1962848 |
| **Antenna connector** | CONN SMA RCPT STR 50 OHM PCB | 733910060 | 1 | **DigiKey** | https://www.digikey.be/product-detail/en/molex/0733910060/WM5543-ND/1465165 |
| **Cable conn. on/off JST-XH ale** | CONN HEADER VERT 2POS 2.5MM | B2B-XH-A-M(LF)(SN) | 1 | **DigiKey** | https://www.digikey.be/product-detail/en/jst-sales-america-inc/B2B-XH-A-M-LF-SN/455-2879-ND/3926507 |
| **power connector 3p male (pcb)** | TERM BLOCK HDR 3POS VERT 3.5MM | 395016003 | 1 | **DigiKey** | https://www.digikey.be/product-detail/en/molex/0395016003/WM25698-ND/2735251 |
| **power connector 3p female** | TERM BLOCK PLUG 3POS 3.5MM | 395038003 | 1 | **DigiKey** | https://www.digikey.be/product-detail/en/molex/0395038003/WM25709-ND/4480336 |
| **in/output connector 7p male** | TERM BLOCK HDR 7POS VERT 3.5MM | 395016007 | 1 | **DigiKey** | https://www.digikey.be/product-detail/en/molex/0395016007/WM25702-ND/2735255 |

| | | | | | |
|---|---|---|---|---|---|
| **in/output connector 7p female** | TERM BLOCK PLUG 7POS 3.5MM | 395037007 | 1 | **DigiKey** | https://www.digikey.be/product-detail/en/molex/0395037007/WM25519-ND/4480323 |
| **conn headers female 12p** | CONN HDR 12POS 0.1 TIN PCB | PPTC121LFBN-RC | 2 | **DigiKey** | https://www.digikey.be/product-detail/en/sullins-connector-solutions/PPTC121LFBN-RC/S6100-ND/807231 |
| **plug-in IC sockets 4x2p** | CONN IC DIP SOCKET 8POS TIN | A 08-LC-TT | 2 | **DigiKey** | https://www.digikey.be/product-detail/en/assmann-wsw-components/A-08-LC-TT/AE9986-ND/821740 |
| **plug-in IC sockets 7x2p** | CONN IC DIP SOCKET 14POS TIN | A 14-LC-TT | 1 | **DigiKey** | https://www.digikey.be/product-detail/en/assmann-wsw-components/A-14-LC-TT/AE9989-ND/821743 |
| **EEPROM** | 512 Kbit, 64K x 8bit, Serial I2C (2-Wire), 400 kHz, DIP, 8 Pins | 24LC512-I/P | 1 | **Farnell** | https://be.farnell.com/microchip/24lc512-i-p/serial-eeprom-512kbit-400khz-dip/dp/9758020?CMP=GRHB-OCTOPART |
| **LoRaWAN module** | RN2483A-I/RM104 or RN2483A-I/RM105 | RN2483A-I/RM104 | 1 | **Farnell** | https://be.farnell.com/microchip/rn2483a-i-rm104/transceiver-module-300kbps-870mhz/dp/2920841?st=rn2483 |
| **Coating spray** | PLASTIK 70 conformal coating | 2646050 | / | **Farnell** | https://be.farnell.com/kontakt-chemie/plastik-70/conformal-coating-aerosol-200ml/dp/2646050 |
| **Battery** | Li-ion 3500mah, 18650, flat top | 12429 | 1 | **VaBo** | http://www.vabo.be/ned/ned_main.htm |
| **Silicone** | Transparant sanitary Kit Handson 310 ml | 604449 | / | **Gamma** | https://www.gamma.nl/assortiment/handson-sanitairkit-transparant-310-ml/p/B604449 |
| **Tie wraps (100 pcs.)** | 200x2,5 mm 100p, Handson | 110695 | / | **Gamma** | https://www.gamma.nl/assortiment/handson-bindband-zwart-2-5-x100-mm-100-stuks/p/B110695 |
| **Solder tin** | 0.7 mm tin/copper 50 gram, Griffon | 607125 | / | **Gamma** | https://www.gamma.nl/assortiment/griffon-printsoldeer-0-7-mm-tin-koper-50-gram/p/B607125 |
| **PCB** | Printed Circuit Board | **/** | 1 | **JLCPCB** | https://jlcpcb.com/ |
| **3D Pint** | PLA print using | **/** | 1 | **FabLab** | https://fablab-leuven.be/?lang=en |
| **wire** | Silicon flexible wire, 7.5m 26AWG | KW-2371 | / | **KIWI electronics** | https://www.kiwi-electronics.nl/kw-2371?search=Siliconen%20Draad%20Soepel&description=true |
| **12 pin header** | 40 pin header strip | KW-2424 | 0.66 | **KIWI electronics** | https://www.kiwi-electronics.nl/40-pin-header-strip?search=header%20pins&description=true |
| **quick conn. on/off tab (10 pcs)** | Tabs till 4.8mm, 2.54mm JST-XH - male | KW-2310 | 0.1 | **KIWI electronics** | https://www.kiwi-electronics.nl/kw-2310 |
| **Plastic containers** | IN: 56x36x50 mm, OUT: 60x40x52(LxBxH), with lid | 31185 | 0.1 | **Bodemschat** | https://www.bodemschat.nl/en/plastic-box-60-x-40-x-52-mm.html |
| **Battery Module** | 5V Micro USB 18650 Lithium Battery Charger,UPS module Step up | **/** | 1 | **AliExpress** | https://nl.aliexpress.com/store/4706110?spm=a2g0o.detail.1000007.1.165664d57AGZiY |

#### 10.2 Electrical Scheme

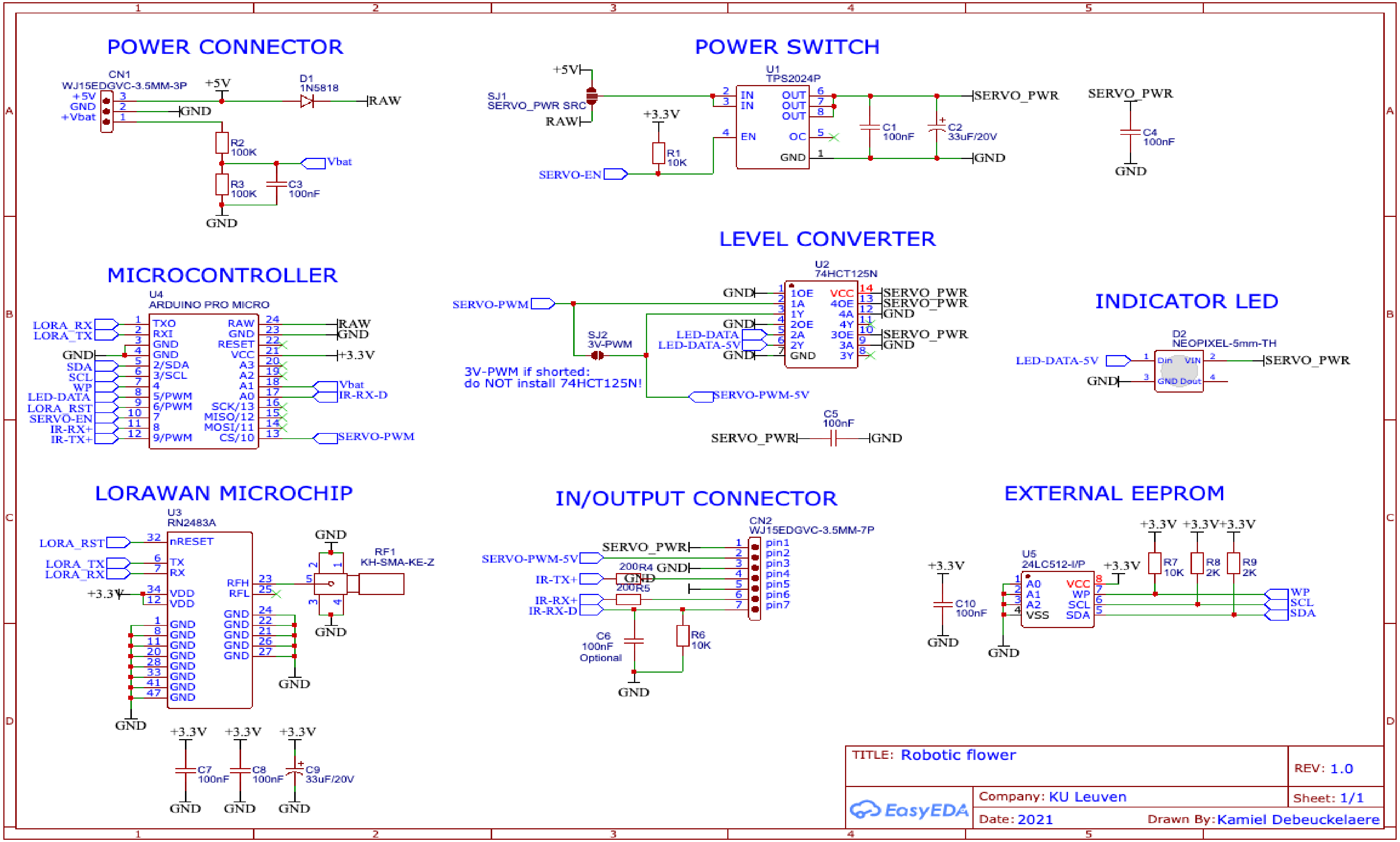
